# MDiNE: A model to estimate differential co-occurrence networks in microbiome studies

**DOI:** 10.1101/544122

**Authors:** Kevin McGregor, Aurélie Labbe, Celia MT Greenwood

## Abstract

**Motivation:** The human microbiota is the collection of microorganisms colonizing the human body, and plays an integral part in human health. A growing trend in microbiome analysis is to construct a network to estimate the co-occurrence patterns among taxa though precision matrices. Existing methods do not facilitate investigation into how these networks change with respect to covariates.

**Results:** We propose a new model called Microbiome Differential Network Estimation `(MDiNE)` to estimate network changes with respect to a binary covariate. The counts of individual taxa in the samples are modelled through a multinomial distribution whose probabilities depend on a latent Gaussian random variable. A sparse precision matrix over all the latent terms determines the co-occurrence network among taxa.

The model fit is obtained and evaluated using Hamiltonian Monte Carlo methods. The performance of our model is evaluated through an extensive simulation study, and is shown to outperform existing methods in terms of estimation of network parameters. We also demonstrate an application of the model to estimate changes in the intestinal microbial network topology with respect to Crohn’s disease.

**Availability and Implementation:** `MDiNE` is implemented in a freely available R package: https://github.com/kevinmcgregor/mdine.

**Supplementary information:** A file containing supplemental material has been submitted with this manuscript.

## 1 Introduction

The human microbiota refers to the microorganisms residing on and inside the human body. These organisms form symbiotic relationships with their host and consequently have a direct effect on human metabolism and host-environment interaction (Turnbaugh et al., 2007). Additionally, the microbiota form ecological communities influenced by body site, metabolic diversity, and competition for resources (Chu et al., 2017; Hibbing et al., 2010). In order to perform any kind of analysis of microbiome data, microbes are classified based on genetic similarity into units called operational taxonomic units (OTUs). By mapping these OTUs to a reference genome, they can be further grouped into taxonomic classifications such as species, family, or phylum. Technologies such as 16S ribosomal RNA sequencing allow researchers to get a snapshot of the taxonomic makeup of microbiome samples in the form of counts of each taxonomic group within each sample.

The co-occurrence patterns among taxa in the human gut microbiome are driven by metabolic interactions and competition for resources (Levy and Borenstein, 2013). However, the development of statistical methodology for the analysis of 16S sequencing data, in addition to other metagenomic sequencing platforms, has largely focused on the relative abundances and diversity of the various taxa found in microbiome samples. Differences in microbial composition have been linked to type I diabetes, obesity, and stunted growth (Brugman et al., 2006; Gough et al., 2015; Turnbaugh et al., 2009). While it is evident that these methods have proven effective in capturing information about changes in microbial composition, they do not directly address the question of relationships between the taxa in a community. Co-occurrence patterns can be represented through the use of networks, where the nodes correspond to taxa and the edges define the correlation patterns between the taxa. Statistical approaches to modeling taxa co-occurrence networks have the potential to capture important community characteristics that would otherwise go undetected in abundance or diversity based analyses. There is also value in estimating the difference between two co-occurrence networks, as it is then possible to investigate how external factors are related to changes in the network structure of the microbiome.

A convenient way to model networks is through precision matrix estimation, as the pattern of zeros occurring in the precision matrix can be used to construct a network. Given a set of *J* random variables, pairwise covariances form a *J* × *J* covariance matrix Σ. The precision matrix is defined as the matrix inverse of the covariance matrix, i.e. Σ^-1^ Drton et al. (2007). The advantage of using the precision matrix lies in the off-diagonal elements, which can be transformed to partial correlations between pairs of variables. That is, the partial correlation *r*_*jj*′_ between variable *j* and variable *j*′, conditional on all other variables can be calculated as:

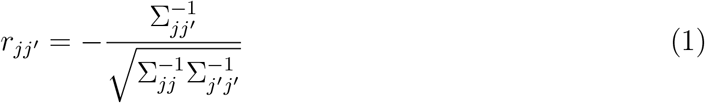

(Kuismin and Sillanpää, 2017). If *r*_*jj*′_ = 0, then variable *j* is uncorrelated with variable *j*′ given all the other variables. As a result, the precision matrix gives an idea of how the variables are directly related in the sense that it removes the possibility that an observed association only exists through correlation with one or more other variables.

In 16S sequencing analysis, nodes in the network are assigned to the microbial taxa and edges correspond to non-zero entries in the precision matrix. In the high-dimensional setting, when the number of taxa is large relative to the sample size, sparsity (or an abundance of zero entries) in the precision matrix must be assumed in order to obtain stable estimates of the precision matrix elements (Chi and Lange, 2014). Furthermore, if the number of taxa exceeds the sample size, then the covariance matrix is singular. The estimation of sparse precision matrices in the Gaussian graphical setting is a well studied problem. Graphical LASSO (GLASSO) was proposed as an efficient solution to the problem, and remains a standard in the domain of sparse precision matrix estimation (Friedman et al., 2008). Nonetheless, several characteristics of 16S sequencing data inhibit the use of Gaussian graphical models. Firstly, since the data are in the form of counts, often with many zeros, the assumption of normality does not hold. More importantly, due to the nature of the sequencing process, the data are compositional; that is, inherent features of the sequencing process lead to varying read depths over samples which can bias the observed correlations or similarity metrics. Proper inference can only be performed on the abundance of a taxon relative to one or more of the other taxa (Gloor et al., 2017).

Several approaches have been developed to estimate networks in microbiome data. `SparCC` and `CCLasso` directly model correlations between taxa in a non-parametric manner (Fang et al., 2015; Friedman and Alm, 2012). `MInt` incorporates a Poisson-Multivariate Normal model which estimates a precision matrix as well as covariate effects on taxon abundances (Biswas et al., 2016). Sparse Inverse Covariance Estimation for Ecological Association Inference `(SPIEC-EASI)` runs GLASSO on the centered log-ratio transformed counts (Kurtz et al., 2015). One common element of each of these methods is that they model a single co-occurrence network among the samples. Yet, there has been interest in estimating separate networks according to a condition, such as a treatment or a disease of interest, to see how the network structure is affected (Bajaj et al., 2012; Gough et al., 2015; Mahana et al., 2016; Ruiz et al., 2017).

Models to compare precision matrices exist in other contexts. The Joint Graphical LASSO was proposed as a solution in the case of Gaussian data (Danaher et al., 2014). Similarly, a non-parametric method for differential network estimation was proposed by Zhao et al. (2014). In this algorithm, the difference in precision matrices is modeled directly, whereas the precision matrix within each group is not explicitly estimated. To our knowledge, there is no existing differential network estimation procedure for microbiome data based on precision matrix estimation. Currently, a network estimation algorithm must be run separately within each group of interest, effectively reducing the sample size used to model each network.

In this paper we introduce a new model, called Microbiome Differential Network Estimation `(MDiNE)`, capable of estimating separate taxa co-occurrence networks for groups defined by a binary variable. The method can also recover the differences between each network, as well as provide interval estimates for each parameter. Furthermore, estimates of how covariates, including the binary variable of interest, associate with the relative abundance of the taxa can also be obtained. We perform a simulation study to compare the performance of MDiNE against two precision matrix-based methods: `SPIEC-EASI` and `MInt`. Additionally, we illustrate the performance of MDiNE through an application on 16S sequencing data from Crohn’s disease and control samples.

## 2 Methods

We have chosen a model that is based on the multinomial distribution, which allows us to address the compositional nature of the data. The Dirichlet-Multinomial distribution has been proposed in this context (Chen and Li, 2013; Holmes et al., 2012), and remains a natural choice given that the Dirichlet distribution is a conjugate prior for the multinomial distribution. Because of varying read depths over samples, covariances should not be calculated on the raw taxa counts. The most obvious way to deal with the problem is to normalize by the total number of counts in each sample, so that the abundance of each taxon *j* ∈ {1, …, *J* + 1} is expressed as a proportion *p*_*j*_. However, this introduces a sum constraint in each sample, such that 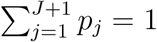. Any attempt at modelling the covariance structure on a proportion would lead to a bias towards negative covariances (van den Boogaart and Tolosana-Delgado, 2013). Calculating covariances on compositional data also does not preserve subcompositional coherence, a property that guarantees that inference on a subset of taxa does not depend on whether or not other taxa are included in the analysis (Aitchison, 1994).

In 2013 Xia et al. introduced a logistic normal multinomial model capable of estimating associations between taxa abundances and covariates that addresses some of the aforementioned challenges (Xia et al., 2013). Define *p* = (*p*_1_, …, *p*_*J*_, *p*_*J*+1_) as a vector of proportions of *J* + 1 different taxa in a sample. The logistic normal multinomial model implicitly models the *additive log-ratio transformation* of *p*, which is defined as:

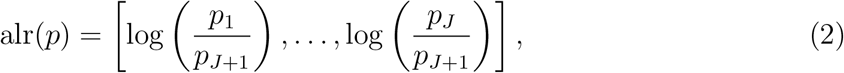

assuming each *p*_*j*_ > 0. This transformation does indeed preserve subcompositional coherence. Multivariate normality is assumed for the transformed counts, meaning that the precision matrix is explicitly defined as a parameter in the model. We therefore pursue an extension to the model posited by Xia et al by including two precision matrices, as well as a penalization scheme to select network edges.

In contrast, the method `SPIEC-EASI` directly computes the centered log-ratios of the vector of *observed* taxa counts *y* = (*y*_1_, …, *y*_*J*_, *y*_*J*+1_) as such:

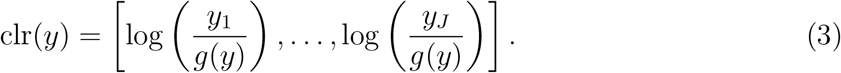

where *g*(*p*) is the geometric mean of the values in *p*

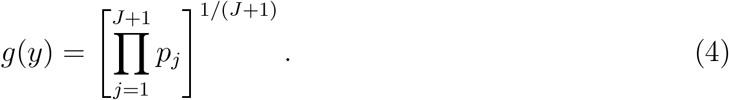

From there, a graphical estimation procedure such as GLASSO is applied to the transformed data to obtain network estimates. Yet, there is typically a large number of zero counts in 16S sequencing data, which cannot be handled in the log-ratio transformation. An arbitrary positive pseudo-count must be added before taking the transformation. The multinomial logistic model, however, can handle zero counts without resorting to the addition of a pseudo-count.

The statistical theory for penalized covariance matrices in a logistic multinomial normal model is complex. We choose to define the model in a Bayesian framework, and obtain the model fit using Markov Chain Monte Carlo (MCMC) methods. The priors placed on the model parameters control the level of parsimony applied in parameter estimation. Details of the MCMC methods used can be seen in Subsection 2.4.

### 2.1 Model

Among *N* individuals we consider *J* taxa for inclusion in the network analysis, and the sum of the counts of the remaining taxa are placed into a reference group *J* + 1. The number of covariates in the model is *K*. The counts of the taxa are contained in the matrix **Y**_*N*×(*J*+1)_. The total number of reads for individual *i* is denoted as 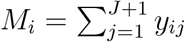. Let *Z* = (*z*_1_, …, *z*_*N*_) be a vector containing the binary covariate over which the taxa co-occurrence networks are expected to vary. **X** = (***1***, *X*_1_, …, *X*_*K*_)_*N*×(*K*+1)_ is the design matrix, and **X**_*i·*_ represents row *i* of the design matrix containing individual *i*’s covariates. Our method estimates how each of the covariates in the design matrix relates to taxon abundance. The vector *Z*, which is of primary interest for the network changes, could also be included in the design matrix.

Assume the true proportions of the *J* +1 taxa in individual *i*’s microbiome is *p*_*i·*_ = (*p*_*i*1_, …, *p*_*i*(*J*+1)_) such that 0 < *p*_*ij*_ < 1 for *j* ∈ {1, …, *J* + 1} and 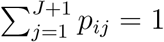. Then the vector of counts for individual *i*, denoted by **Y**_*i·*_, can be modeled as a multinomial distribution:

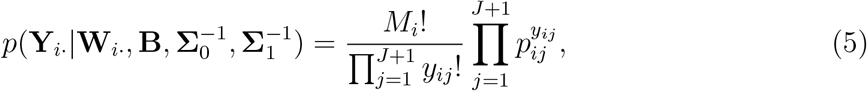

where the proportions are parameterized through the additive log-ratio transformation with respect to category *J* + 1 as seen in Equation 2:

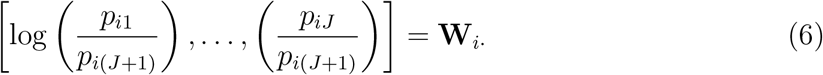

for *j* ∈ {1, …, *J*}. Here we model the row vector of log-ratio transformed proportions for individual *i*, **W**_*i·*_, as a multivariate normal distribution whose parameters depend on the observed covariates **X** and *Z*:

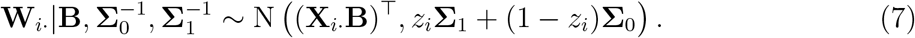

That is, the variance structure depends on the individual’s binary covariate *z*_*i*_. The corresponding precision matrices 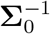 and 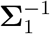 therefore define the co-occurrence network structure of the *J* taxa of interest within each group defined by *Z*. The parameters in the matrix **B**_(*K*+1)×*J*_ define how each of the *K* covariates in **X** relate to the abundances of the taxa relative to the reference category *J* + 1.

Equivalently, the inverse transformation for the additive log-ratio transformed data is expressed as:

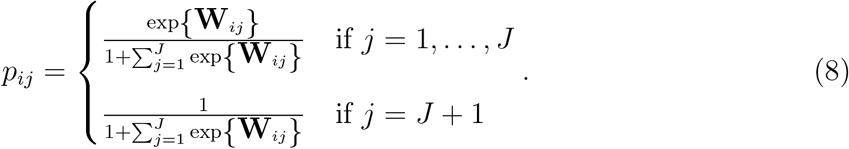

Given that we are implementing a Bayesian framework, non-informative (high-variance) normal priors are assumed for each element of **B**. The full model is summarized in Equation 14.

The potentially large number of parameters in 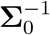 and 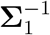 relative to the sample size *N* necessitates a procedure to shrink the off-diagonal elements of these two matrices. We now outline a prior-based shrinkage strategy in the model.

### 2.2 Inducing sparsity

To solve the problem of unstable estimates of precision matrices resulting from inadequate sample size (as mentioned in the introduction), we must apply a penalization scheme to the elements of 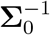 and 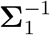 In a Bayesian framework, strategic priors can be used on the model parameters to shrink estimates towards zero. In our model, we apply a Laplace prior centered around zero and with inverse scale parameter λ to induce sparsity in the off-diagonal elements of the two precision matrices. This is the Bayesian equivalent to adding an *L*_1_ penalty term to the log-likelihood in the maximum likelihood framework, where the following maximization would take place:

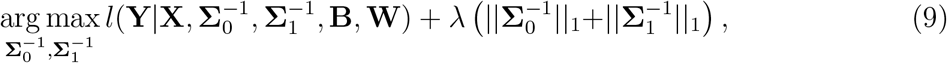

where *l* represents the log-likelihood of the hierarchical model as defined in Section 2.1. ||*·*||_1_ denotes the sum of the absolute values of the matrix elements of 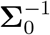 or 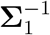.

As the diagonal elements are constrained to the set of positive real numbers, an exponential prior with rate parameter λ*/*2 is used for these elements instead (Khondker et al., 2013; Wang, 2012). Let 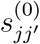 and 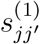 represent the value in row *j* and column *j*′ of the matrices 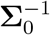 and 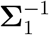, respectively. Then the priors for the elements of 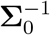 are:

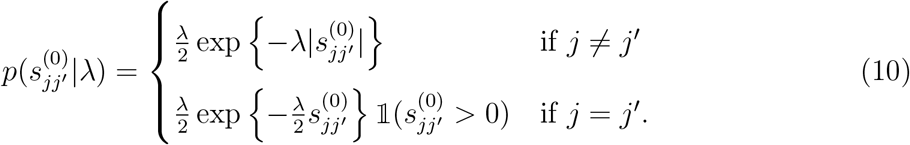

The priors for the elements of 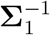 are defined likewise with the same λ. This single value of λ will then control the amount of parameter shrinkage in the two precision matrices. The current parameterization implies that a higher value of λ will correspond to smaller elements in absolute value; that is, a precision matrix with off-diagonal elements closer to zero. The strategy for choosing this penalization parameter is outlined in the next subsection.

### 2.3 Choice of λ

In the MCMC framework, the value of the penalization parameter λ can be specified as a hyperparameter in the model along with its own prior distribution. In the Bayesian Graphical Lasso, a Gamma prior is applied to the Laplace penalization parameter (Wang, 2012), since this leads to its full conditional itself following a Gamma distribution. In our model, we specify an exponential prior on λ, as it is a special case of the Gamma distribution, but only includes a single hyperparameter. In order to avoid choosing an arbitrary hyperparameter in the exponential prior, we use an empirical Bayes-like procedure that is similar to what is outlined in (Biswas et al., 2016).

We start by estimating an initial multinomial logistic regression model using the counts **Y** as the outcome and the covariates **X** as predictors, which gives an initial estimate 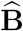. From there, we obtain the residuals of the model (on the log-ratio scale), 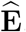 and calculate the empirical covariance matrices separately for residuals of samples with *z*_*i*_ = 0 and *z*_*i*_ = 1. To obtain initial, naïve estimates 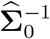 and 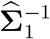, we take the generalized inverses of the empirical covariance matrices. We can plug in our initial estimates 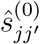 and 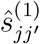 into 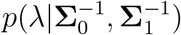 and maximize with respect to λ to obtain an initial estimate:

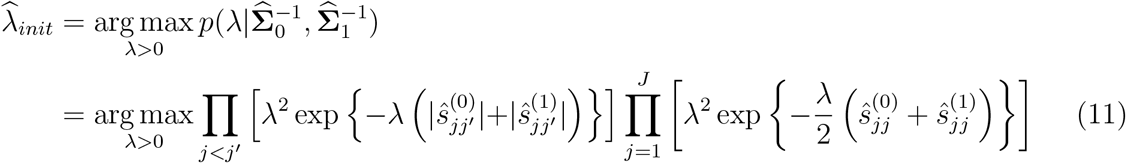

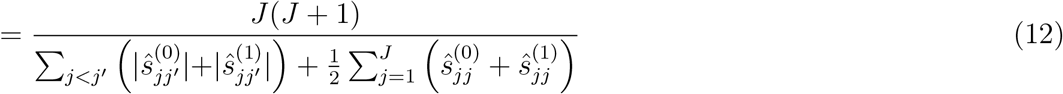

In the full model, we specify an exponential prior on λ having expected value equal to the initial estimate 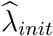 so that λ is sampled over a reasonable region:

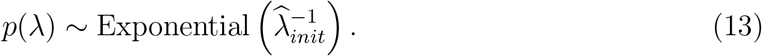

More details on the choice of the parameter λ and how it can be estimated can be seen in Section 1 of the Supplementary File. In summary, the full model can then be written as:

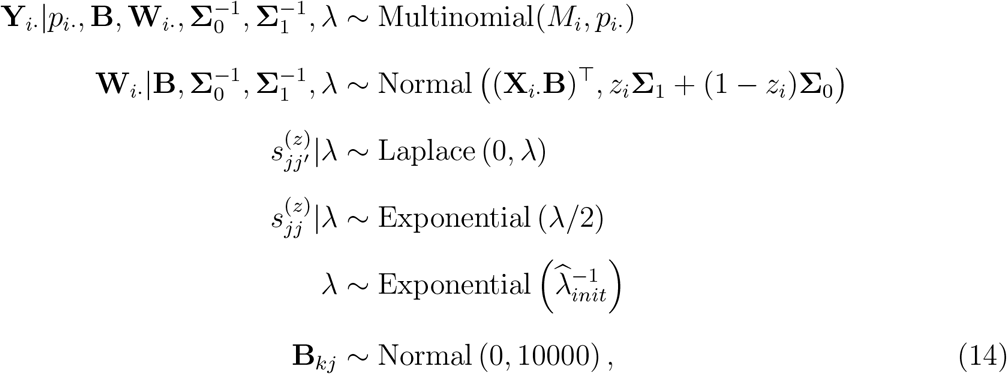

for each *i* ∈ {1, …, *N*}, *j* ∈ {1, …, *J*}, *j*′ ∈ {1, …, *j* - 1}, *k* ∈ {1, …, *K* + 1}, and *z* ∈ {0, 1}.

The expression for the joint posterior distribution 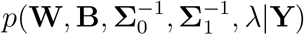 can be seen in Section 2 of the Supplementary File.

### 2.4 Parameter estimation

The model is fit through Hamiltonian Monte Carlo (HMC) sampling facilitated by the Stan language (Carpenter et al., 2017). One particular advantage of HMC is its efficiency in the case of correlated parameters, which can be a significant hurdle in other Monte Carlo simulation strategies (Rannala, 2002). To ensure positive definiteness of 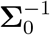 and 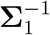, we parameterize them through their Cholesky decompositions, so that 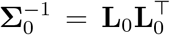 and 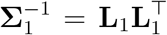. Note that Stan also allows matrices to be declared as covariance matrices, hence preserving positive definiteness during the HMC sampling process. Estimates for each parameter are obtained by calculating the posterior mean from the obtained MCMC samples. Each model was run for 1000 samples in 4 chains, with the first half of samples taken as warm-up. Convergence was verified through trace plots and calculation of the potential scale reduction factor.

### 2.5 Simulation design

A simulation study was undertaken to evaluate the performance of `MDiNE` against that of `MInt` and `SPIEC-EASI`. To ensure realistic count data, we based the simulation parameters on data coming from the American Gut Project, which is a publicly available, crowdfunded human microbiome database http://humanfoodproject.com/americangut. The zero-inflated negative binomial (ZINB) model was previously shown to be a good choice for simulating realistic microbiome count data (Kurtz et al., 2015). Therefore, we chose to simulate data through the ZINB model in a way that replicated the distributions of the most common fecal microbial families found in the American Gut data.

The data generating process involved the “Normal to Anything” method (Cario and Nelson, 1997; Kurtz et al., 2015), where correlated taxa vectors were simulated from a multivariate normal distribution and were transformed to a ZINB model based on the American Gut data. ZINB parameters were estimated separately for individuals with asthma (310 subjects) and without asthma (3193 subjects). The parameters for subjects with asthma were used to simulate data for subjects in the *z*_*i*_ = 1 group and the parameters for subjects without asthma were used for the *z*_*i*_ = 0 group. Details on the American Gut data simulation procedure can be seen in Section 3 of the Supplementary File.

We ran twelve simulation scenarios, with differing sample sizes and numbers of taxa: *N* ∈ {50, 100, 500, 750} and *J* ∈ {5, 10, 25}. We ran 100 replications for each scenario.

We also performed a simulation where data were generated directly from the model assumed in `MDiNE`. This simulation allowed investigation of parameter bias and precision for the precision matrix elements in `MDiNE`, which cannot be obtained through the first data generation mechanism. Details of this additional simulation study can be found in the Section 4 of the Supplementary File.

### 2.6 Metrics of comparison

Since the model assumptions are substantially different between `MDiNE` and the other two network methods, the values in the estimated precision matrices cannot be directly compared. Consequently, our primary metrics of performance are sensitivity and specificity of network edge detection and a measure of overall network structure. In particular, we consider the area under the receiver operating curve (AUC) for detection of edges. Even for these metrics, the comparison of methods is tricky. Although the parameter λ controls the amount of shrinkage in `MDiNE`, none of the off-diagonal elements of 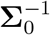 or 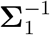 can be set exactly to zero, in contrast to `MInt` and `SPIEC-EASI`. As such, we considered an edge to exist in the `MDiNE` estimates if its credible interval did not contain zero. The use of interval estimates to direct variable selection has precedent in Bayesian penalization procedures (Khondker et al., 2013; Li et al., 2010; Park and Casella, 2008). We note, however, that such an approach should only be used to detect potentially important links between taxa. Setting elements of a precision matrix to zero based on interval estimates does not necessarily preserve positive definiteness.

It is also valuable to compare networks based on measures that capture the overall network structure. *Weighted natural connectivity* is a measure of structural robustness in that it measures the extent to which the connectivity of the network is vulnerable to edge deletion (Xiao-Ke et al., 2013). Details on its calculation can be seen in Section 5 of the Supplementary File.

### 2.7 Data application

We performed a network analysis on a dataset from Gevers et al. (2014). The data consisted of publicly available 16S sequencing measures from a multi-cohort study of new-onset Crohn’s disease. To avoid potential problems arising from mixing samples from different cohorts, we only analyzed samples from the Risk Stratification and Identification of Immunogenetic and Microbial Markers of Rapid Disease Progression in Children with Crohn’s Disease (RISK) study https://clinicaltrials.gov/ct2/show/NCT00790543. The participants in the RISK cohort were all 18 years of age or younger and were recruited from 28 centres in Canada and the USA. Cases were subjects with newly diagnosed Crohn’s disease and controls were subjects presenting with non-inflammatory gastrointestinal conditions.

Samples from mucosal tissue biopsies of the terminal ileum were available for 314 Crohns patients and 192 controls. The analysis focused on the family taxonomic level, with 15 families included in the analysis. We examined network differences between Crohn’s samples and controls (i.e. *z*_*i*_ = 1 for Crohn’s patients, and *z*_*i*_ = 0 for controls). The Crohn’s disease status was also included in the model matrix **X**, in addition to age, sex, and antibiotic use. The dataset included preprocessed `.biom` files, and OTU tables and taxonomic information were extracted using the `phyloseq` package in `R` (McMurdie and Holmes, 2013). Results from the data application can be seen in Section 3.3.

## 3 Results

### 3.1 Simulation results

Here we present the results of the American gut dataset based simulation study which employed the performance metrics outlined in Subsection 2.6, and compare the results of `MDiNE` to those of `SPIEC-EASI` and `MInt`. In Figures S2 and S3, we show that the simulated data reasonably replicates the distribution of the observed counts in the American Gut study data.

First, we examined the AUC achieved within each estimation method. Figure 1 shows AUC distributions over 100 simulation scenarios. The results are shown separately for groups *z* = 0 and *z* = 1, and are further split up by number of taxa (*J*) and sample size (*N*). It is evident that `MDiNE` generally outperforms the other methods in terms of AUC across all scenarios. It also appears for `MDiNE` that there is an improvement in AUC as the sample size *N* increases. There is also a considerable amount of variability across datasets in AUC among all methods in the *J* = 5 scenario.

**Figure 1:**
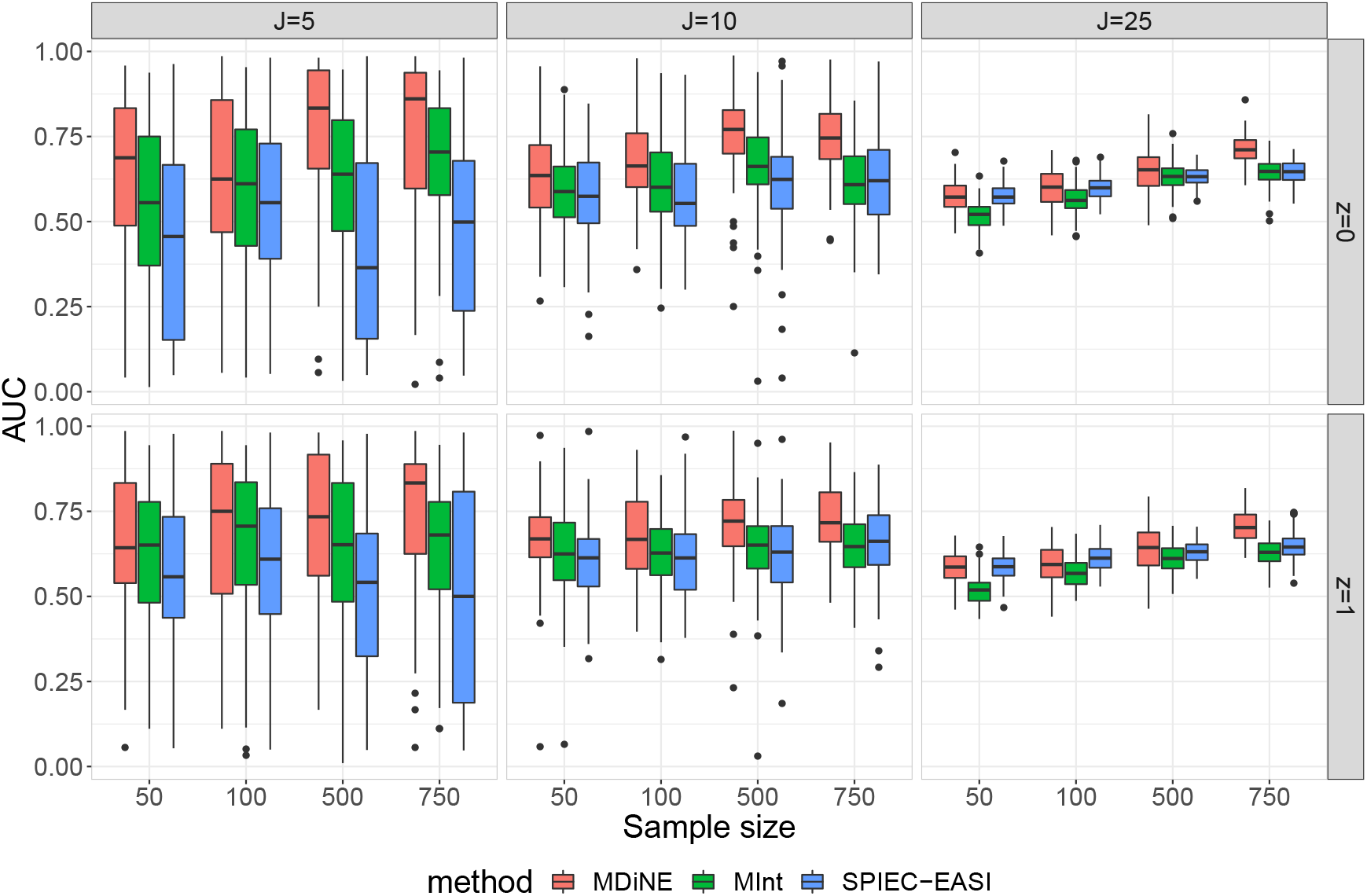
The AUC for network edge detection in each of `MDiNE`, `MInt`, and `SPIEC-EASI` based on simulated data. The results vary over the number of taxa included in the network (*J*), the sample size (*N*), and the two precision matrices corresponding to groups *z* = 0 and *z* = 1.

One important observation is the reduced accuracy of edge detection in all methods in the *J* = 25 scenario. Even with the large sample size *N* = 750, the AUC is generally less than 0.75. This result may not be surprising considering there are 25*×*(25+1)*/*2 = 325 parameters to estimate within each precision matrix, for a total of 650 parameters, in addition to the covariate effects in **B**. Even if many elements of the matrix are zero, there are a large number of parameters to estimate relative to the sample size. Since all three methods use some kind of selection or shrinkage procedure, there is generally a bias towards zero in the estimates of the precision matrix elements.

In order to assess overall structure of the estimated networks, we calculated the weighted natural connectivity (based on partial correlations), as defined in Equation 12 in the Supplementary File, and compared to the natural connectivity in each true simulated partial correlation matrix. Figure 2 shows the distribution of the logarithm of the squared error (square of the estimated minus simulated values) of the natural connectivity over the simulation replications.

**Figure 2:**
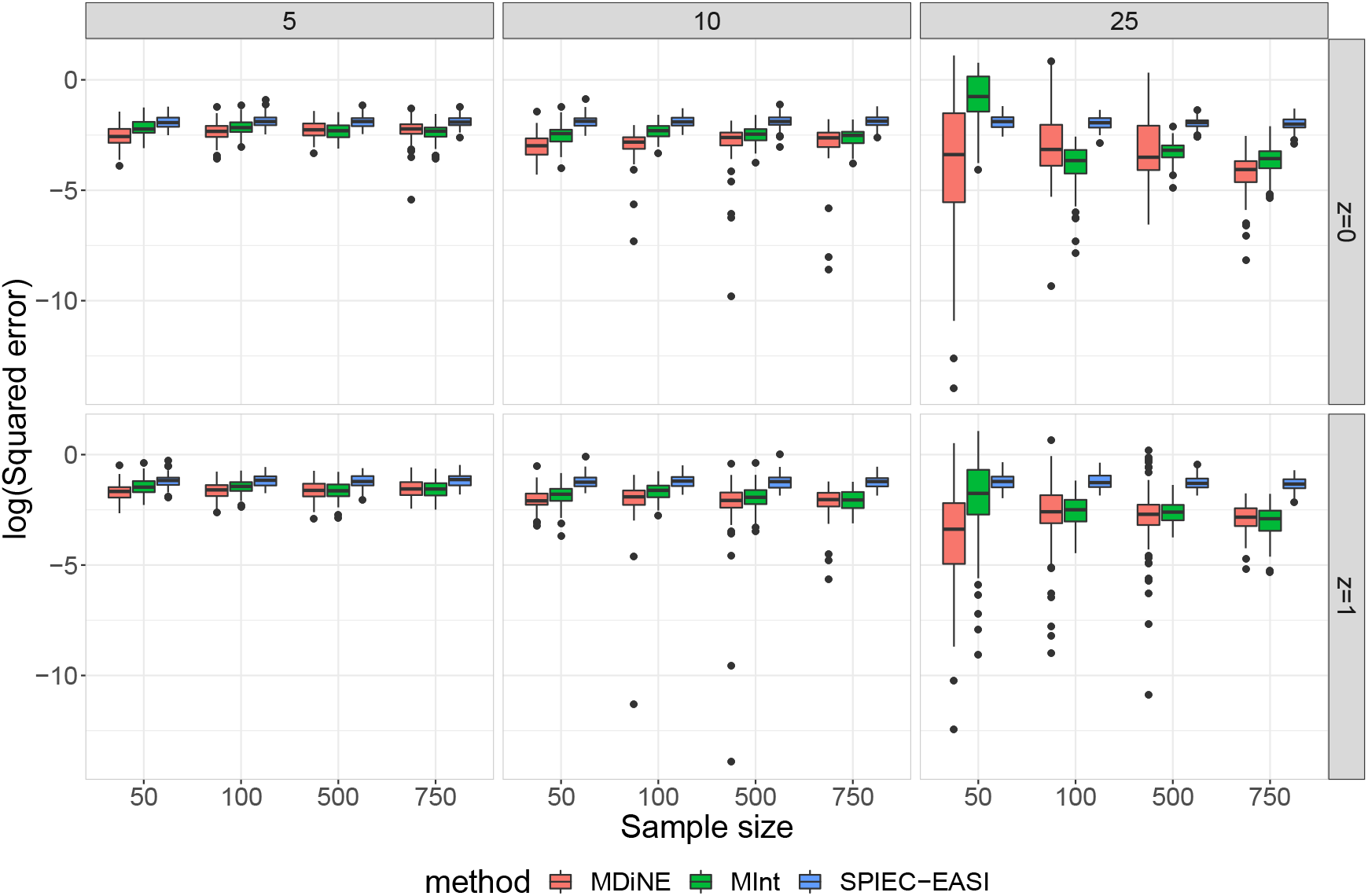
Log squared error of the weighted natural connectivity in each of `MDiNE`, `MInt`, and `SPIEC-EASI` based on simulated data. The results vary over the number of taxa included in the network (*J*), the sample size (*N*), and the two precision matrices corresponding to groups *z* = 0 and *z* = 1.

In general, `MDiNE` outperformed the other methods in terms of estimating natural connectivity. Interestingly, there was not much improvement in estimation accuracy as sample size increased. There was also an increase in the variance of estimation accuracy for `MDiNE` in the *J* = 25 case. Other methods also demonstrated increased variability in this measure of network accuracy, but less so than `MDiNE`.

### 3.2 Testing parameter estimation in MDiNE

#### 3.2.1 Estimation accuracy

Next, we examined the results of the simulation where data were generated under the model assumed in `MDiNE`. Figure S4 shows the squared errors averaged over the elements of 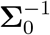 and 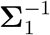. As expected, there was a considerable improvement in estimation accuracy as sample size increased. For a fixed sample size the errors increased with respect to the number of taxa included in the analysis. We also plotted the squared errors for the difference in the precision matrices 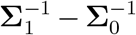 in Figure S5, and observed a similar result. The same pattern was also present, though to a lesser extent, in the abundance effects **B** (Figure S6). The estimation accuracy for **B** generally improved over sample size.

We also performed an analysis on credible interval coverage, which can be viewed in Section 6.2 the Supplementary File. The analysis showed that coverage greatly depended on the sample size relative to the number of parameters in the model.

#### 3.2.2 Effect of *λ*

To demonstrate the shrinkage effects of the Laplace and exponential priors on the elements of 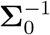 and 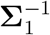, we plotted the estimated values of these elements over values of the penalization parameter *λ* (Figure 3). It is clear that the value of *λ* had a profound impact on the magnitudes of the estimated parameters in both precision matrices. For values greater than 2^10^ the estimates for all parameters were very close to zero (though never exactly equal to zero). We note that specifying an unnecessarily large value for the penalization parameter could cause issues with the HMC sampler and hence lead to unstable parameter estimates.

**Figure 3:**
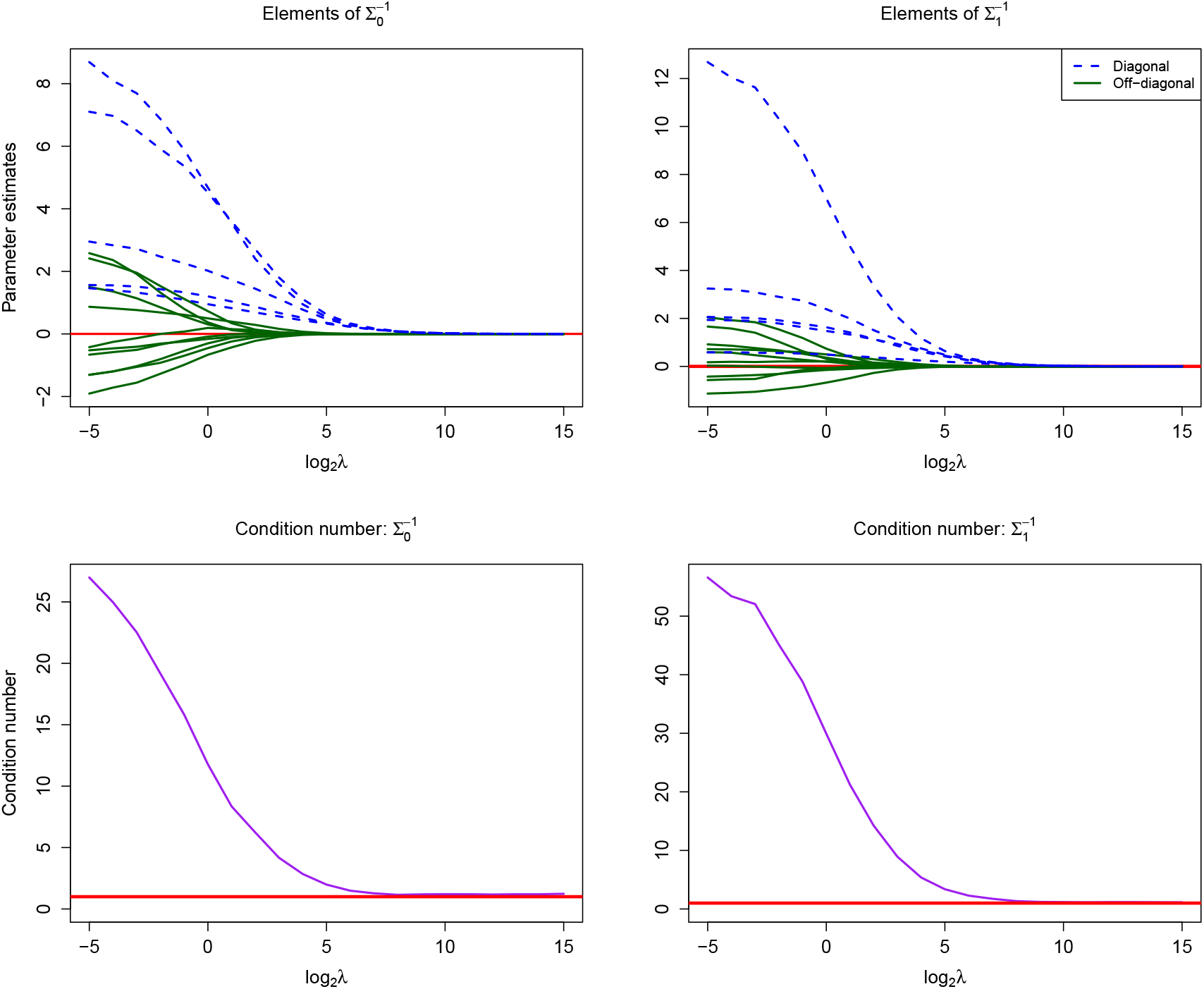
(Top) The `MDiNE` coefficient shrinkage paths for elements of 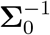 and 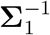 over values of *λ*. (Bottom) The condition number of each matrix over the values of *λ*. The data came from the *N* = 50, *J* = 5 scenario. Note the horizontal axes are on a base-2 logarithmic scale.

The impact of *λ* was also evaluated through the condition numbers of 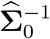 and 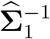. The condition number of a positive-definite matrix is calculated as the ratio of the largest and smallest eigenvalues of the matrix (Won et al., 2013). A large condition number implies that a covariance matrix is ill-conditioned, leading to numerical instability in inversion. The condition number for the estimated values of the two matrices over different values of *λ* are shown in Figure 3. For very small values of *λ* the condition number was large. The condition number steadily decreased for larger values of *λ*, clearly demonstrating that the penalization parameter was able to prevent against an ill-conditioned precision matrix when its size was large relative to the sample size. The distributions of the estimated values of *λ* over the 100 simulation replications are shown in Figure S1. As expected, higher values of *λ* were generally estimated for higher numbers of taxa included in the network analysis.

### 3.3 Data application results

We applied `MDiNE` to the Crohn’s disease dataset described in Section 2.7. We chose the top 15 represented families based on the mean count over all individuals to include in the network analysis. The counts for all remaining families were merged into the reference category. The estimated family networks for Crohn’s and control samples are shown in Figure 4. It appears that there were more positive co-occurrences in the control samples, whereas there were some additional negative co-occurrences in the Crohn’s samples. There were also important differences in the relative abundances of Bacteroidaceae, Porphyromonadacea, and Lachnospiraceae.

**Figure 4:**
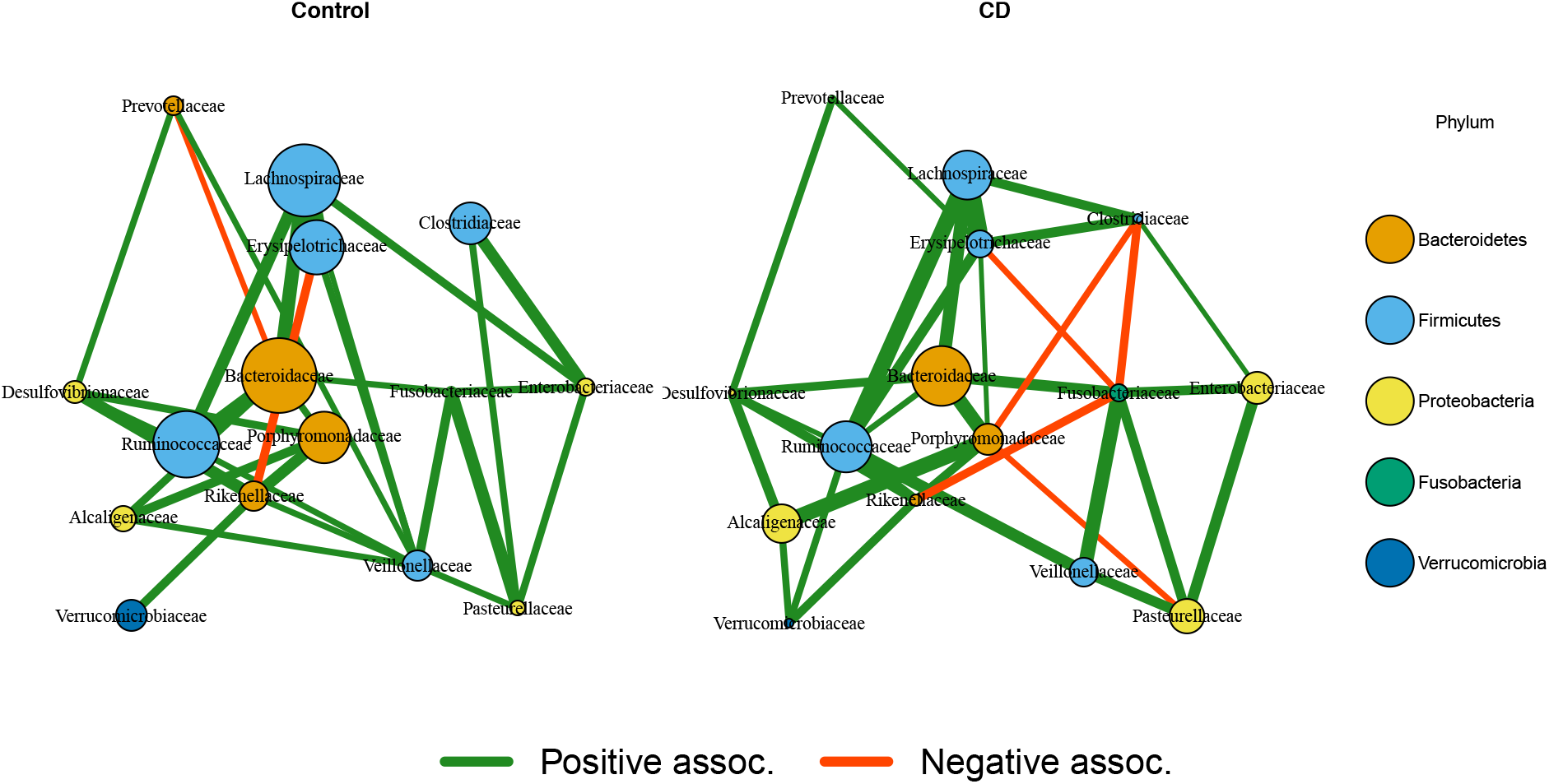
Estimated co-occurrence from `MDiNE` at the family level in control (left) and Crohn’s (right) samples. An edge is displayed if its 90% credible interval does not contain zero. The size of a node represents its relative abundance. The placement of the nodes is based on multidimensional scaling of the precision matrix in the Crohn’s samples.

An important advantage of `MDiNE` over existing methods is the ability to detect significant changes in family co-occurrences with respect to Crohn’s status. Figure 5 characterizes the differences between the Crohn’s and control networks. Only associations that significantly differ between the two networks, based on 90% credible intervals, were included in the figure. Based on this criterion, only five family-family co-occurrence associations differed significantly between the Crohn’s cases and controls: Lachnospiraceae and Enterobacteriaceae; Lachnospiraceae and Erysipelotrichaceae; Erysipelotrichaceae and Fusobacteriaceae; Erysipelotrichaceae and Veillonellaceae; and Porphyromonadaceae and Pasteurel-Next, we compared the estimated networks between `MDiNE`, `SPIEC-EASI`, and `MInt`. Since `SPIEC-EASI` cannot handle covariates, we estimated the networks without adjusting for age, sex, or antibiotic status. Results can be seen in Section 7.1 of the Supplementary File.

**Figure 5:**
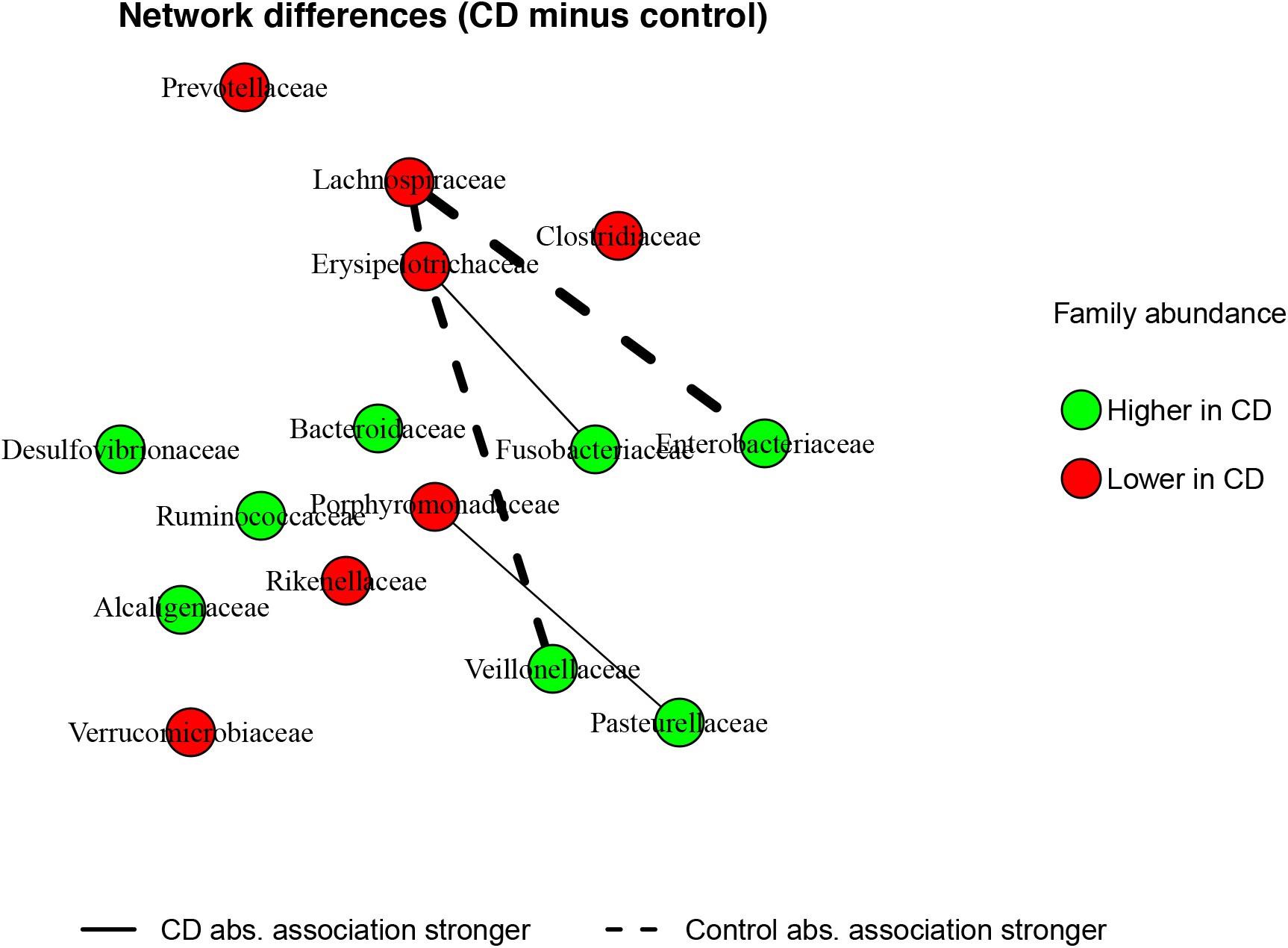
Network differences between Crohn’s and control samples. An edge is shown when the co-occurrence between a pair of taxa differs significantly between the two groups based on the 90% credible interval. Colours show whether family abundance is higher in Crohn’s or in control samples.

Figure S11 contains the estimated networks in Crohn’s cases and control samples for each of the estimation methods. It is evident that the edge density differed greatly between the methods, with `MInt` resulting in a very edge-dense network, and `SPIEC-EASI` resulting in a much more sparse network. `SPIEC-EASI` also appears to have mostly identified negative taxon co-occurrences, whereas both `MDiNE` and `MInt` uncovered both positive and negative co-occurrences.

A natural question to ask is: to what extent do the three methods agree with respect to network edges? To test this we plotted the concordance between the methods in a Venn diagram, shown in Figure S12. There appears to be much more concordance between `MDiNE` and `MInt` with respect to network edges, as they agreed on 30 edges in the control samples, and 34 edges in the Crohn’s samples. There were only 6 edges common to both `MDiNE` and `SPIEC-EASI`. These 6 edges were identified in all three methods.

The convergence of `MDiNE` was verified in Section 7.2 of the Supplementary File. Traceplots and examination of the potential scale reduction factor confirmed proper mixing of chains and adequate convergence. These results are displayed in Figures S13-16.

This analysis has shown that there are important differences in co-occurrences between several highly abundant families in the intestinal microbiota of Crohn’s and control samples. Significance of such differences would not have been detected using previous network estimation methods.

## 4 Discussion

We have introduced a new microbiome network estimation model that generally outperformed two existing precision matrix-based methods, both in terms of estimating individual taxon associations, as well as overall network structure. More importantly, we have defined our model in a way that differences between the associations in two networks defined by a binary variable could be directly estimated. As the model parameters were estimated through MCMC sampling, interval estimates for all model parameters were easily obtained. Though we presented an example from the 16S sequencing platform, `MDiNE` could be used on any count-based platform, such as whole genome sequencing.

Attention must be drawn to the more broad issue of sample size in network inference. If the desire is to perform inference on individual co-occurrences in the network, then it should be noted that applying a network estimation method with built-in regularization will tend to under estimate the number of true associations if the sample size is not sufficiently large. Even for a modest number of taxa (say 25), a relatively large sample size is required to ensure adequate credible interval coverage, due to the rapidly increasing number of parameters to estimate with increasing network size. We believe this fact has not been properly explored in the literature, and extra study of this problem would be beneficial.

An important limitation of `MDiNE` is its running time. Though MCMC sampling has its advantages, the computational burden is always a factor. Regardless, `MDiNE` benefits from a Bayesian implementation because interval estimates can be obtained for both the network edges as well the changes in taxon-taxon associations with respect to the binary variable *Z*. Existing methods would have to rely on bootstrap sampling schemes or edge stability measures in order to determine the importance of associations between taxa. These additional procedures would greatly increase their computational times.

For the moment the MDiNE model only applies to a single binary covariate defining two networks. In future work we plan to extend the method to estimate how the networks change with respect to a continuous covariate. This is a challenging problem, as the estimated precision matrix must be positive-definite for every possible value of the covariate. Some additional structure will need to be assumed in the parameterization of the precision matrix to satisfy this constraint.

## 5 Conclusion

We have developed a powerful new precision matrix-based network estimation tool called `MDiNE` which facilitates the comparison of networks defined by a binary covariate. Furthermore, interval estimates can be obtained both for network parameters as well as the difference between networks, which is not currently possible in existing microbiome network estimation methods.

## 6 Software

We have developed an R package called `mdine`. The package is currently available on Github https://github.com/kevinmcgregor/mdine, and will soon be submitted to the Comprehensive R Archive Network (CRAN).

