## Supplementary File for "MDiNE: A model to estimate differential co-occurrence networks in microbiome studies"

#### 1 Details on choosing the penalization parameter $\lambda$

Here we explain the details of the choice of the penalization parameter $\lambda$, which was summarized in the methods section of the main manuscript. We fit an initial multinomial logistic regression model such that, for each sample $i$:

$$\mathbf{Y}_{i\cdot} \sim \text{Multinomial}(M_i, p_{i\cdot}), \tag{1}$$

with each $p_{ij}$, $j \in \{1, \ldots, J+1\}$ parameterized as:

$$p_{ij} = \begin{cases} \frac{\exp\{\mathbf{X}_{i\cdot}\mathbf{B}_{\cdot j}\}}{\sum_{j=1}^{J}\exp\{\mathbf{X}_{i\cdot}\mathbf{B}_{\cdot j}\}+1} & \text{if } j = 1, \ldots, J \\ \frac{1}{\sum_{j=1}^{J}\exp\{\mathbf{X}_{i\cdot}\mathbf{B}_{\cdot j}\}+1} & \text{if } j = J+1 \end{cases}. \tag{2}$$

This initial model is fit using maximum likelihood estimation, which has been implemented for multinomial logistic regression in the `R` package `nnet` (Ripley et al., 2016). The estimated value for $\mathbf{B}$ is denoted as $\widehat{\mathbf{B}}$. The residuals of the model for sample $i$ and taxon $j$ (on the additive log-ratio scale) are computed as:

$$\widehat{\mathbf{E}}_{ij} = \log\left(\frac{y_{ij}+1}{y_{i(J+1)}+1}\right) - \mathbf{X}_{i\cdot}\widehat{\mathbf{B}}\cdot j, \tag{3}$$

where a pseudo-count of 1 is added to each element of $\mathbf{Y}$ as zero counts cannot be accommodated in the log-ratio transformation. Assume, without loss of generality, that the rows of $\widehat{\mathbf{E}}$ can be rearranged as:

$$\widehat{\mathbf{E}}' = \begin{bmatrix} \widehat{\mathbf{E}}^{(0)} \\ \widehat{\mathbf{E}}^{(1)} \end{bmatrix},$$

where $\widehat{\mathbf{E}}^{(0)}$ and $\widehat{\mathbf{E}}^{(1)}$ contain the rows of $\widehat{\mathbf{E}}$ for samples corresponding to groups $z_i = 0$ and $z_i = 1$, respectively.

Initial estimates of the precision matrices $\mathbf{\Sigma}_0^{-1}$ and $\mathbf{\Sigma}_1^{-1}$, denoted as $\widehat{\mathbf{S}}^{(0)}$ and $\widehat{\mathbf{S}}^{(1)}$, can be obtained by calculating the empirical precision matrices obtained from the model residuals:

$$\widehat{\mathbf{S}}^{(z)} = \left(\widehat{\text{cov}}\left[\widehat{\mathbf{E}}^{(z)}\right]\right)^{+}, \tag{4}$$

for each $z \in \{0, 1\}$. The operator $\mathbf{A}^{+}$ denotes the Moore-Penrose inverse of the matrix $\mathbf{A}$ which is used to handle the possibility that $\mathbf{A}$ is not invertible. If $\mathbf{A}$ is indeed invertible, then $\mathbf{A}^{+} = \mathbf{A}^{-1}$.

Then let $\hat{s}_{jj'}^{(0)} = \widehat{\mathbf{S}}_{jj'}^{(0)}$ and $\hat{s}_{jj'}^{(1)} = \widehat{\mathbf{S}}_{jj'}^{(1)}$ be initial estimates of $s_{jj'}^{(0)}$ and $s_{jj'}^{(1)}$, respectively. To pick a reasonable hyperparameter for the exponential prior on $\lambda$, we begin by maximizing $p(\lambda|\mathbf{\Sigma}_0^{-1}, \mathbf{\Sigma}_1^{-1})$ with respect to $\lambda$ after plugging in the initial estimates calculated above. The goal is to calculate an initial guess $\widehat{\lambda}_{init}$ as follows:

$$\widehat{\lambda}_{init} = \underset{\lambda > 0}{\arg\max}\, p(\lambda|\widehat{\mathbf{\Sigma}}_0^{-1}, \widehat{\mathbf{\Sigma}}_1^{-1}). \tag{5}$$

To begin, we can write the conditional distribution of $\lambda$ as:

$$\begin{aligned} p(\lambda|\mathbf{\Sigma}_0^{-1}, \mathbf{\Sigma}_1^{-1}) &= \prod_{j<j'} \left[\lambda \exp\left\{-\lambda|s_{jj'}^{(0)}|\right\} \times \lambda \exp\left\{-\lambda|s_{jj'}^{(1)}|\right\}\right] \prod_{j=1}^{J} \left[\lambda \exp\left\{-\frac{\lambda}{2} s_{jj}^{(0)}\right\} \times \lambda \exp\left\{-\frac{\lambda}{2} s_{jj}^{(1)}\right\}\right] \\ &= \prod_{j<j'} \left[\lambda^2 \exp\left\{-\lambda\left(|s_{jj'}^{(0)}|+|s_{jj'}^{(1)}|\right)\right\}\right] \prod_{j=1}^{J} \left[\lambda^2 \exp\left\{-\frac{\lambda}{2}\left(s_{jj}^{(0)} + s_{jj}^{(1)}\right)\right\}\right]. \end{aligned} \tag{6}$$

After taking the logarithm, we obtain:

$$\log(p(\lambda|\mathbf{\Sigma}_0^{-1}, \mathbf{\Sigma}_1^{-1})) = \sum_{j<j'} \left[2\log(\lambda) - \lambda\left(|s_{jj'}^{(0)}|+|s_{jj'}^{(1)}|\right)\right] + \sum_{j=1}^{J} \left[2\log(\lambda) - \frac{\lambda}{2}\left(s_{jj}^{(0)} + s_{jj}^{(1)}\right)\right] \tag{7}$$

$$= (J^2 + J)\log(\lambda) - \lambda \sum_{j<j'} \left(|s_{jj'}^{(0)}|+|s_{jj'}^{(1)}|\right) - \frac{\lambda}{2} \sum_{j=1}^{J} \left(s_{jj}^{(0)} + s_{jj}^{(1)}\right), \tag{8}$$

Differentiating with respect to $\lambda$ gives:

$$\frac{\partial \log(p(\lambda|\mathbf{\Sigma}_0^{-1}, \mathbf{\Sigma}_1^{-1}))}{\partial\lambda} = \frac{J^2+J}{\lambda} - \sum_{j<j'} \left(|s_{jj'}^{(0)}|+|s_{jj'}^{(1)}|\right) - \frac{1}{2}\sum_{j=1}^{J}\left(s_{jj}^{(0)} + s_{jj}^{(1)}\right). \tag{9}$$

We set $\frac{\partial \log(p(\lambda|\mathbf{\Sigma}_0^{-1}, \mathbf{\Sigma}_1^{-1}))}{\partial\lambda} = 0$ and solve for $\lambda$. At this time, we can plug in the intial estimates in $\widehat{\mathbf{S}}^{(0)}$ and $\widehat{\mathbf{S}}^{(1)}$ to get the initial estimate of $\lambda$ as:

$$\widehat{\lambda}_{init} = \frac{J(J+1)}{\sum_{j<j'}\left(|\hat{s}_{jj'}^{(0)}|+|\hat{s}_{jj'}^{(1)}|\right) + \frac{1}{2}\sum_{j=1}^{J}\left(\hat{s}_{jj}^{(0)} + \hat{s}_{jj}^{(1)}\right)}. \tag{10}$$

This initial estimate of is used to form the rate hyperparameter in the exponential prior of $\lambda$. We set the mean of this distribution to be equal to $\widehat{\lambda}_{init}$, and so the prior for $\lambda$ is set to be:

$$p(\lambda) \sim \text{Exponential}\left(\widehat{\lambda}_{init}^{-1}\right). \tag{11}$$

To give an idea of the estimated values of $\lambda$ from `MDiNE`, we show in Figure S1 the distributions of $\lambda$ over the American Gut simulation replications.

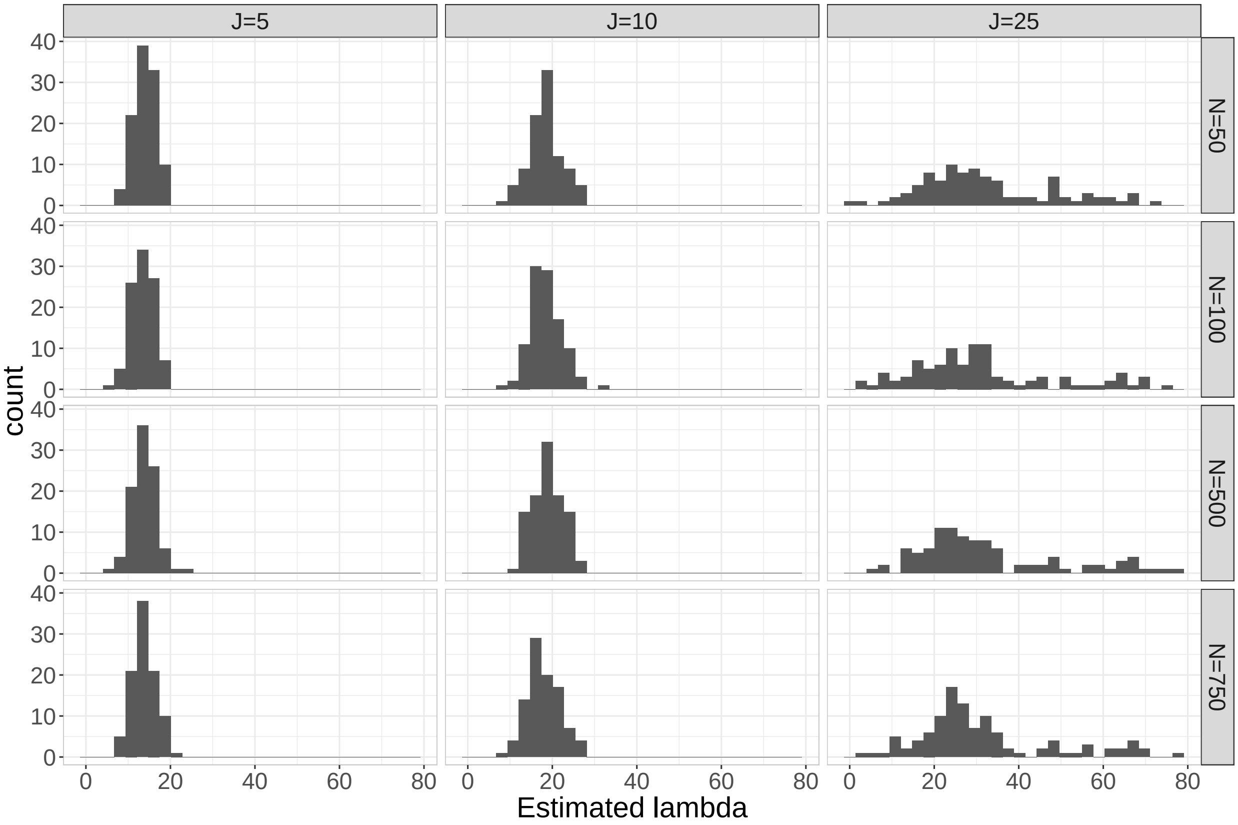


Figure S1: The estimated values of $\lambda$ over the 100 simulation replications in the American Gut simulation study.

### 2 Joint posterior distribution

The joint posterior distribution for the full model outlined in the main manuscript can be written as:

$$
\begin{aligned}
p(\mathbf{W}, \mathbf{B}, \boldsymbol{\Sigma}_0^{-1}, \boldsymbol{\Sigma}_1^{-1}, \lambda|\mathbf{Y}) &\propto p(\mathbf{Y}|\mathbf{W}, \mathbf{B}, \boldsymbol{\Sigma}_0^{-1}, \boldsymbol{\Sigma}_1^{-1}, \lambda) p(\mathbf{W}|\boldsymbol{\Sigma}_0^{-1}, \boldsymbol{\Sigma}_1^{-1}, \lambda) p(\boldsymbol{\Sigma}_0^{-1}|\lambda) p(\boldsymbol{\Sigma}_1^{-1}|\lambda) p(\mathbf{B}) p(\lambda) \\
&= \prod_{i=1}^{N} \left[ \prod_{j=1}^{J} \left( \frac{\exp\{\mathbf{W}_{ij}\}}{\sum_{j=1}^{J} \exp\{\mathbf{W}_{ij}\} + 1} \right)^{y_{ij}} \right] \left( \frac{1}{\sum_{j=1}^{J} \exp\{\mathbf{W}_{ij}\} + 1} \right)^{y_{i(J+1)}} \\
&\times \prod_{i=1}^{N} |(1 - z_i)\boldsymbol{\Sigma}_0^{-1} + z_i \boldsymbol{\Sigma}_1^{-1}|^{1/2} \exp\left\{ -\frac{1}{2} (\mathbf{W}_{i\cdot} - \mathbf{X}_{i\cdot}\mathbf{B}) \left( (1 - z_i)\boldsymbol{\Sigma}_0^{-1} + z_i \boldsymbol{\Sigma}_1^{-1} \right) (\mathbf{W}_{i\cdot} - \mathbf{X}_{i\cdot}\mathbf{B})^\top \right\} \\
&\times \prod_{j=1}^{J} \lambda^2 \exp\left\{ -\lambda \left( s_{jj}^{(0)} + s_{jj}^{(1)} \right) \right\} \prod_{j'=1}^{j} \lambda^2 \exp\left\{ -\lambda \left( |s_{jj'}^{(0)}| + |s_{jj'}^{(1)}| \right) \right\} \mathbb{1}(\boldsymbol{\Sigma}_0^{-1} \in M^+) \mathbb{1}(\boldsymbol{\Sigma}_1^{-1} \in M^+) \\
&\times \prod_{j=1}^{J} \prod_{k=1}^{K+1} 10000^{-1/2} \exp\left\{ -\frac{1}{2 \times 10000} \mathbf{B}_{kj}^2 \right\} \\
&\times \lambda_{init}^{-1} \exp\left\{ -\lambda_{init}^{-1} \lambda \right\},
\end{aligned}
$$

where $M^+$ denotes the set of positive definite, symmetric matrices. The variance for the normal priors on the elements of $\mathbf{B}$ was arbitrarily set to $10,000$ to induce a high-variance prior on those parameters.

### 3 Simulating data based on American Gut data

As mentioned in the main manuscript, the simulation data were generated based on estimated parameters from the American Gut study (amg). The data simulation procedure is outlined in the main manuscript. Here we outline how the simulation parameters were chosen.

The .biom files were obtained and the R package `phyloseq` (McMurdie and Holmes, 2013) was used to extract count tables at the family taxonomic level. We chose to estimate parameters separately for individuals with asthma, and those without. This allowed the specification of slightly different parameters within the $z = 0$ and $z = 1$ groups in the simulation. After discarding samples where asthma status was missing, there were 310 subjects with asthma and 3193 without asthma.

The 50 most highly represented families, based on the mean count over all subjects, were chosen to form the basis of the simulation study. Counts for remaining taxa were quite small and were ignored. Zero-inflated negative binomial (ZINB) parameters were estimated separately for each taxon. The ZINB distribution for a random variable $Y \in \{0, 1, 2, \dots\}$ is defined by (Fang, 2013):

$$P(Y=y)=\begin{cases}\pi+(1-\pi)\left(1+k\mu\right)^{-1/k} & \text{for } y=0\\ (1-\pi)\frac{\Gamma(y+1/k)}{\Gamma(y+1)\Gamma(1/k)}\frac{(k\mu)^y}{(1+k\mu)^{y+1/k}} & \text{for } y=1,2,\ldots\end{cases},$$

where $\pi$ is the zero-inflation parameter, $\mu$ is the mean of the negative binomial distribution, and $k$ is the dispersion parameter of the negative binomial distribution.

In the final simulation, data were generated for 50 taxa regardless of the value of $J$ in the simulation under consideration. However, any generated taxa in excess of the $J$ taxa considered in a single simulation scenario were combined into a single reference category (taxon $J+1$).

To ensure positive-definiteness of the simulated sparse precision matrices, we first generated a sparse Cholesky factor for each matrix. Parameters in each Cholesky factor were chosen so that the resulting non-zero associations in the precision matrix were moderate to strong. This allowed a good comparison of the edge detection performance of the different methods. The data were simulated as follows:

1. For the $N$ subjects randomly set half to $z_i=0$ and half to be $z_i=1$.
2. Generate two lower-triangular matrices $\mathbf{L}_0$ and $\mathbf{L}_1$:
   - For each lower-triangular element of $\mathbf{L}_0$ sample from $Unif(-1.5,1)$ and for diagonal elements, $Unif(1.5,2.5)$.
   - For each lower-triangular element of $\mathbf{L}_1$ sample from $Unif(-2.5,2.5)$ and for diagonal elements, $Unif(2,4)$.
3. Randomly set lower-triangular elements of $\mathbf{L}_0$ and $\mathbf{L}_1$ to zero with probability 0.85.
4. Create the two precision matrices: $\mathbf{\Sigma}_0^{-1}=\mathbf{L}_0\mathbf{L}_0^\top$ and $\mathbf{\Sigma}_1^{-1}=\mathbf{L}_1\mathbf{L}_1^\top$.
5. Generate two $n\times(J+1)$ matrices from multivariate normal distributions $\mathbf{U}^{(0)}\sim \text{MVN}(0,\mathbf{R}_0)$ and $\mathbf{U}^{(1)}\sim\text{MVN}(0,\mathbf{R}_1)$, where $\mathbf{R}_0$ and $\mathbf{R}_1$ are the correlation matrices corresponding to $\mathbf{\Sigma}_0$ and $\mathbf{\Sigma}_1$, respectively.
6. Plug each column of $\mathbf{U}^{(0)}$ and $\mathbf{U}^{(1)}$ into the standard normal cumulative distribution function $\Phi$ and define:
   - $\mathbf{C}_{\cdot j}^{(0)}=\Phi(\mathbf{U}_{\cdot j}^{(0)})$ and $\mathbf{C}_{\cdot j}^{(1)}=\Phi(\mathbf{U}_{\cdot j}^{(1)})$, for $j=1,\ldots,J+1$.
7. Obtain taxon counts by plugging the columns of $\mathbf{C}^{(0)}$ and $\mathbf{C}^{(1)}$ into the quantile function of the ZINB distribution with parameters specific to taxon $j$, denoted by $G_j^{-1}$:

- $\mathbf{Y}_{ij} = \begin{cases} G_j^{-1}(\mathbf{C}_{ij}^{(0)}) & \text{if } z_i = 0 \\ G_j^{-1}(\mathbf{C}_{ij}^{(1)}) & \text{if } z_i = 1. \end{cases}$

Initially, the ZINB parameters were estimated using maximum-likelihood estimation on the real data. However, attempts to evaluate the three network estimation methods were unsuccessful, as all methods showed consistently poor performance over all simulation scenarios. As this simulation setup did not facilitate a good comparison between methods, we chose to instead fix the dispersion parameter at $k = 0.75$, and estimate the $\mu$ and $\pi$ based on observed mean and zero proportion of each taxon. This led to a much better performance comparison between methods (as seen in the main manuscript), and the distribution of the simulated data reasonably resembled the underlying American Gut data.

To demonstrate this, we first examined the distribution of the read depths of the simulated data and compared to that of the true data. This can be seen in Figure S2. The distribution of total read counts was highly skewed, but the skewness of the simulated data was slightly reduced relative to the real data.

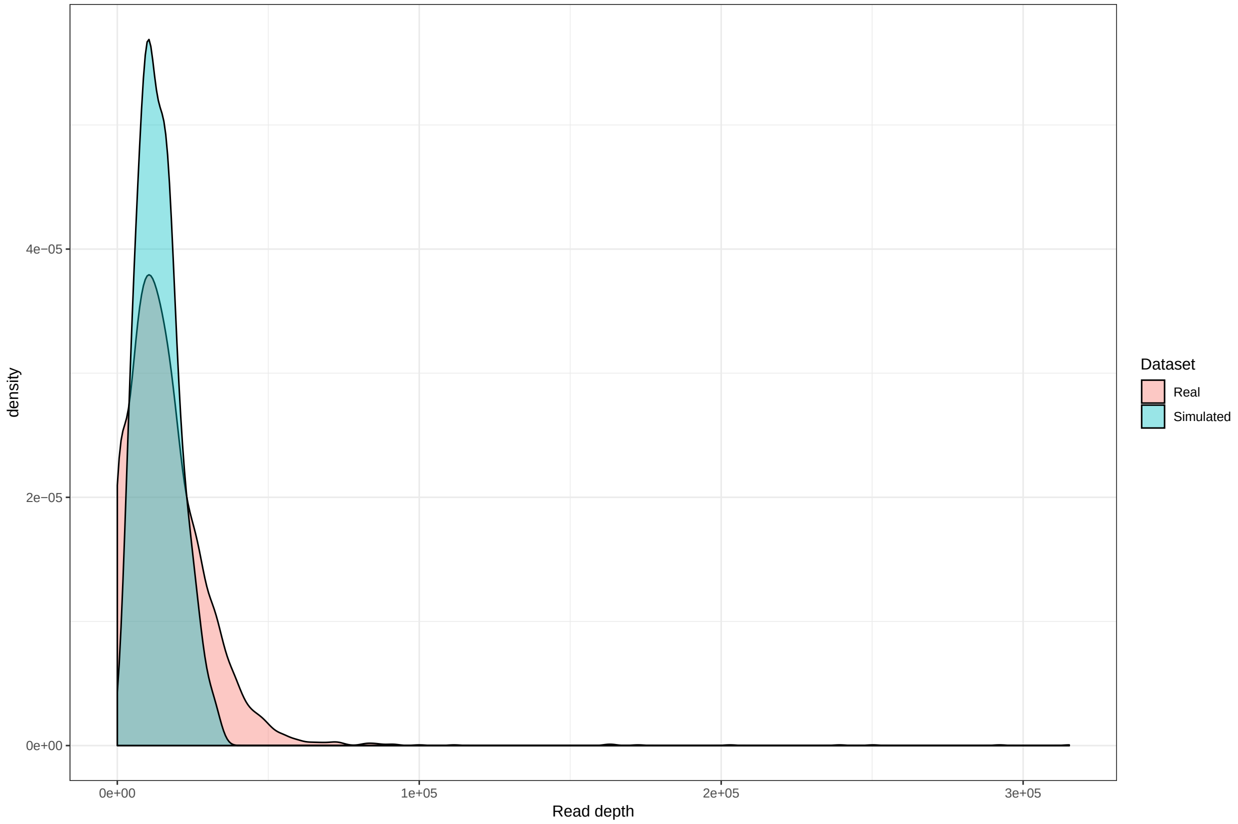


Figure S2: The observed distributions of the read depths of the simulated and real (American Gut) data.

Next we examined how the distribution of each simulated taxon compared with the distribution of the corresponding taxon from the American Gut data. Figure S3 shows QQ-plots to make this comparison. For the most part, the simulated distributions taxa appeared to be reasonably similar to the true distributions. One taxon showed an appreciable deviation from the real data (middle left in the figure). However, this seems to be a reasonable outcome when considering the abnormal distributions of taxon counts usually observed in microbiome data.

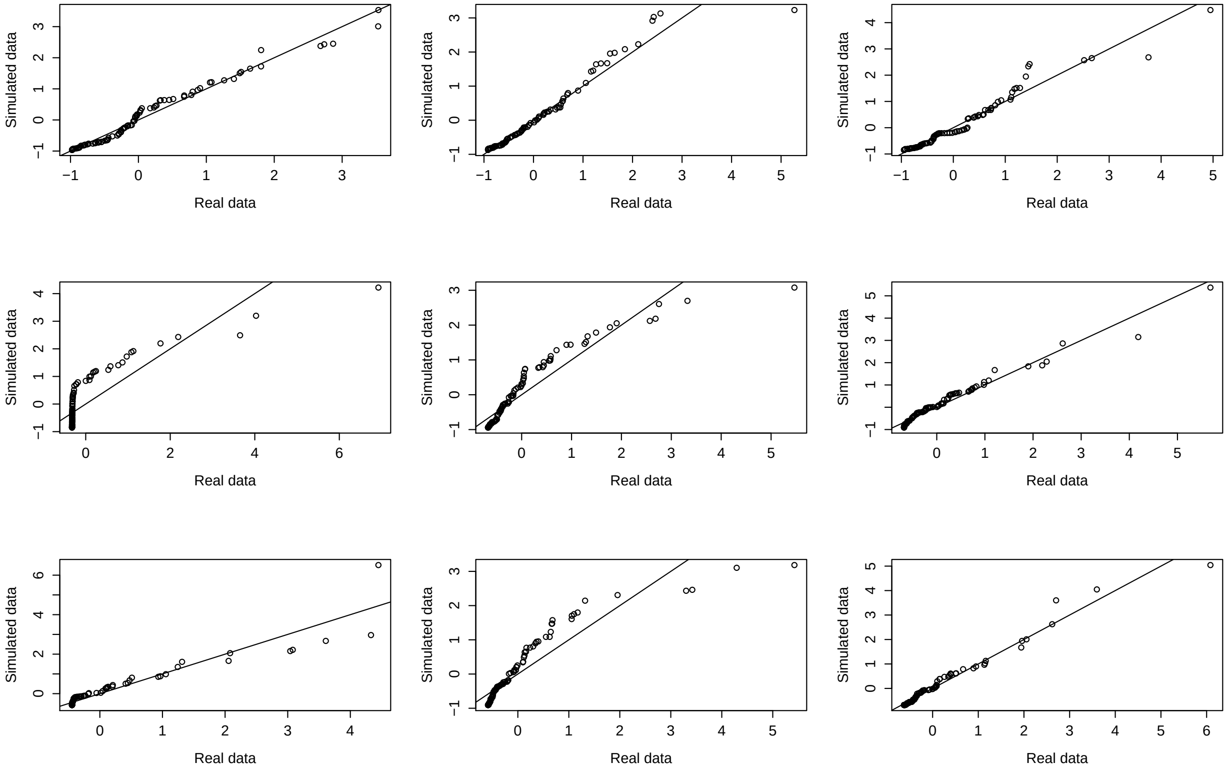


Figure S3: QQ-plots comparing the distribution of the real vs. simulated taxa. The nine most highly represented taxa are shown, one taxon in each plot.

### 4 Simulation based on multinomial model

In addition to the simulation based on the American Gut study, we also performed a simulation based on the model assumed in `MDiNE` to evaluate the performance of parameter estimation within `MDiNE`. Once again simulations were run for sample sizes $N \in \{50, 100, 500, 750\}$, and numbers of taxa $J \in \{5, 10, 25\}$. Here, parameters in the Cholesky decompositions of $\mathbf{\Sigma}_0^{-1}$ and $\mathbf{\Sigma}_1^{-1}$ were chosen to give moderate to strong associations. The distribution of the read depths was estimated from an unpublished dataset containing 16S sequencing data from Rheumatoid Arthritis patients.

1. For the $N$ subjects randomly set half to $z_i = 0$ and half to be $z_i = 1$.

2. Construct the design matrix as $\mathbf{X} = (\mathbf{1}, Z)$

3. Generate two lower-triangular matrices $L_0$ and $L_1$. The parameters were originally chosen based on the rheumatoid arthritis data, but were altered to create stronger associations:

   - For each lower-triangular element of $L_0$ sample from $Unif(-4, 4)$ and for diagonal elements, $Unif(1.5, 2.5)$.

   - For each lower-triangular element of $L_1$ sample from $Unif(-4.5, 4.5)$ and for diagonal elements, $Unif(4, 2)$.

4. Randomly set lower-triangular elements of $L_0$ and $L_1$ to zero with probabilities 0.75 and 0.85, respectively.

5. Create the two precision matrices: $\mathbf{\Sigma}_0^{-1} = L_0 L_0^\top$ and $\mathbf{\Sigma}_1^{-1} = L_1 L_1^\top$.

6. Simulate the fixed effect parameter matrix $\mathbf{B}$ from $N(3, 4)$ for the intercept column, and $N(0, 4)$ for the column corresponding to $Z$.

7. $W \sim N(\mathbf{XB}, z_i \mathbf{\Sigma}_1 + (1 - z_i)\mathbf{\Sigma}_0)$

8. Obtain $p_{ij}$ based on Equation 8 in the main manuscript.

9. Simulate total counts for subject $i$ as: $N_i \sim N(112874, 12163^2)$. These values were the mean and standard error of total reads from the rheumatoid arthritis data.

10. Generate multinomial counts $\mathbf{Y}_{i\cdot} \sim \text{Multinomial}(M_i, p_{i\cdot})$

### 5 Performance metric details

In addition to the AUC metrics defined in the main manuscript, it is also valuable to compare networks based on measures that capture the overall network structure. *Weighted natural connectivity* is a measure of structural robustness in that it measures the extent to which the connectivity of the network is vulnerable to edge deletion (Xiao-Ke et al., 2013). Higher values of the weighted natural connectivity correspond to networks with either more connections, stronger connections, or both. As the precision matrices in each method are estimated through different models, we instead calculated the natural connectivity based on the partial correlation matrices. More precisely, Define the matrix $\mathbf{A}$, such that $\mathbf{a}_{jj'} = |r_{jj'}|$, as the weighted adjacency matrix corresponding to a network defined by scaling the precision matrix $\mathbf{\Sigma}^{-1}$ as seen in Equation 1 of the main manuscript, with diagonal $\mathbf{a}_{jj} = 1$. Let $\gamma_1, \gamma_2, \ldots, \gamma_J$ be the eigenvalues of $\mathbf{A}$. Then we define the weighted natural connectivity to be:

$$\overline{\gamma} = \log\left(\frac{1}{J}\sum_{j=1}^{J} e^{\gamma_j}\right) \tag{12}$$

The resulting weighted natural connectivity of each network estimation method was calculated and compared against the weighted natural connectivity of the true scaled precision matrices.

In the simulation study based on the model assumed in `MDiNE`, we could directly compare the estimated values with the true simulated values of the parameters. For the $\mathbf{B}$ parameter, the absolute errors were calculated for each matrix element were then averaged to obtain a single value for the entire matrix. To compare these average errors over the different simulation replications and scenarios, we divided by the true simulated values to interpret the error as a percentage.

$$\text{PCT-Error}(\mathbf{B}) = \frac{1}{JK}\sum_{j=1}^{J}\sum_{k=1}^{K}\left|\frac{\widehat{\mathbf{B}}_{kj} - \mathbf{B}_{kj}}{\mathbf{B}_{kj}}\right|. \tag{13}$$

As there are zero values in the simulated $\boldsymbol{\Sigma}_0^{-1}$ and $\boldsymbol{\Sigma}_1^{-1}$, the same metric could not be used. Instead, we simply looked at the squared difference between the estimated and simulated parameters, again averaged over all matrix elements.

### 6 Performance of parameter estimation in `MDiNE`

In this section we present additional exploration of the performance of `MDiNE` in terms of parameter estimation and credible interval coverage. All results in this section were obtained from the simulation study outlined in Section 4 of this Supplementary File, where the data are simulated under the model assumed in `MDiNE`.

#### 6.1 Parameter estimation accuracy

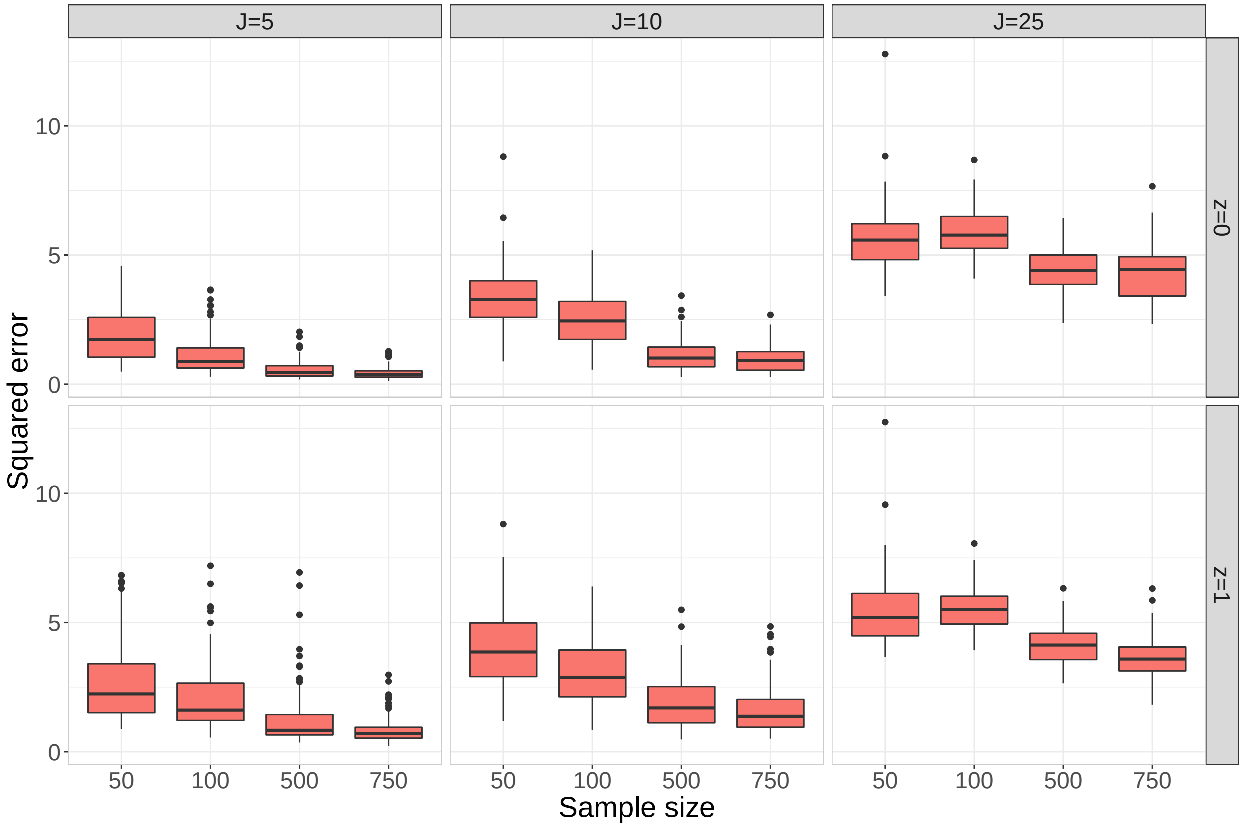


Figure S4: The squared error, $(estimated - true)^2$, averaged over all elements of $\boldsymbol{\Sigma}_0^{-1}$ (top) and $\boldsymbol{\Sigma}_1^{-1}$ (bottom). The boxplots show the distribution over all simulation replications.

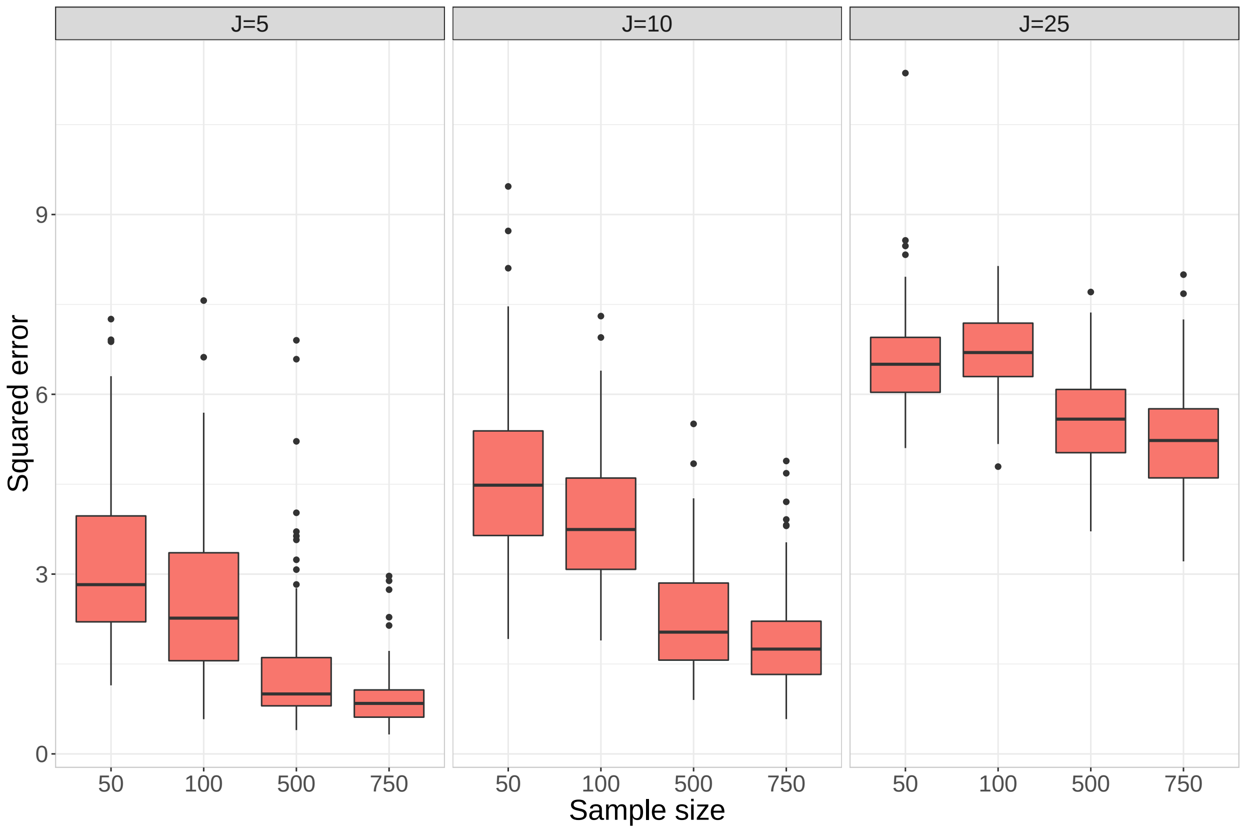


Figure S5: The squared error, $(estimated - true)^2$, averaged over all elements of the difference $\mathbf{\Sigma}_1^{-1} - \mathbf{\Sigma}_0^{-1}$. The boxplots show the distribution over all simulation replications.

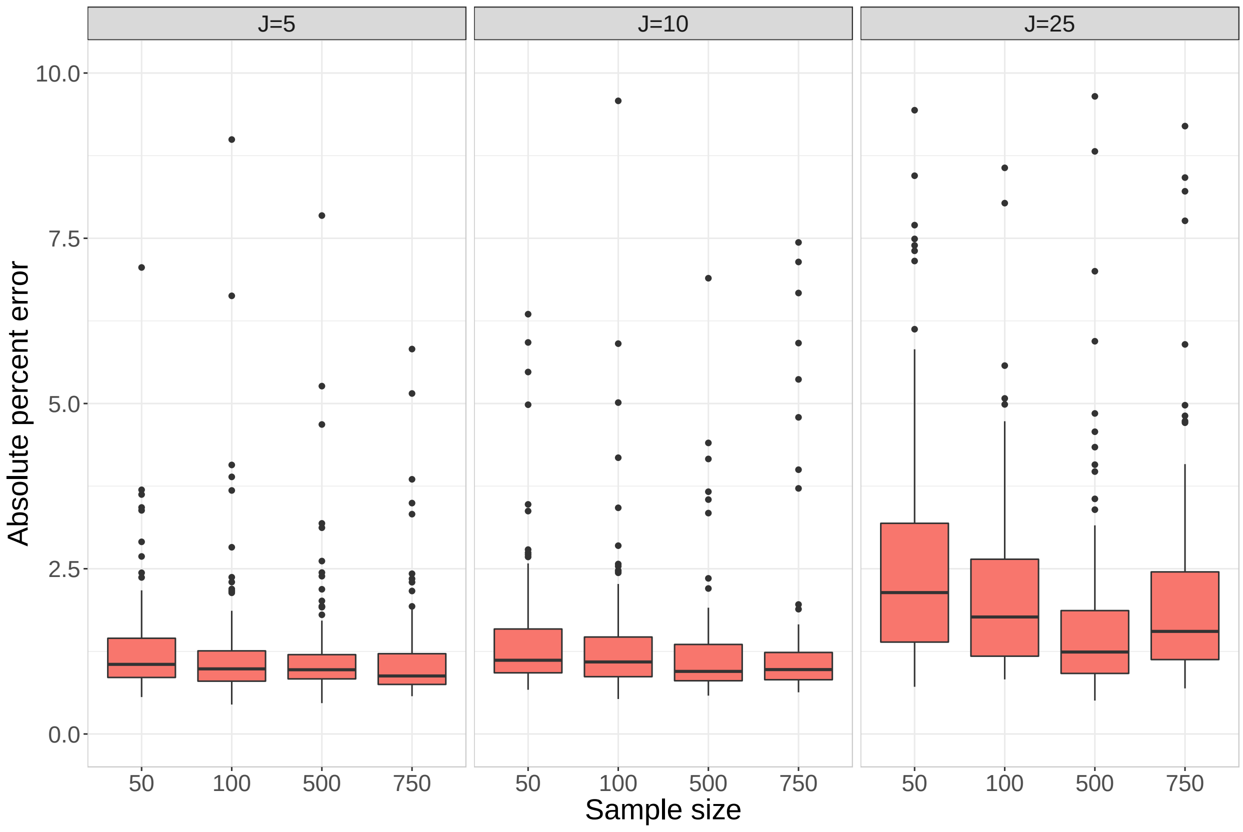


Figure S6: The absolute percent error of the elements in $\mathbf{B}$. For each simulation replication, the absolute percent error between the estimated and simulated values of $\mathbf{B}$ are averaged together. The boxplots show the distribution of this average error over all simulation replications.

#### 6.2 Credible interval coverage

One of the main advantages of `MDiNE` is the ability to extract credible intervals for the parameters in $\boldsymbol{\Sigma}_0^{-1}$, $\boldsymbol{\Sigma}_1^{-1}$, and their difference. We constructed 90% credible intervals for the individual elements of $\boldsymbol{\Sigma}_0^{-1}$ and $\boldsymbol{\Sigma}_1^{-1}$ and estimated the coverage, i.e. the probability that the intervals overlap with the true simulated values. Figure S7 shows the distribution of credible interval coverage (averaged over the elements of each precision matrix). In the $J = 5$ case the coverage appeared to be around 90% for all sample sizes. In the $J = 10$ case the coverage depended on sample size, with coverage approaching the desired 90% with increased sample size. For $J = 25$ coverage was well below the 90% threshold in all sample sizes. The coverage for the $z = 1$ group was generally higher, likely due to the fact that that matrix was simulated to be more sparse.

Figures S8 and S9 show coverage for elements of $\boldsymbol{\Sigma}_0^{-1}$ and $\boldsymbol{\Sigma}_1^{-1}$ for true zero elements and true non-zero elements, respectively. For smaller sample sizes the coverage for true zero elements was greater than the expected 90%. Coverage for true non-zero elements improved drastically with increasing sample size. The result is similar in Figure S10, where the coverage for the difference $\mathbf{\Sigma}_1^{-1} - \mathbf{\Sigma}_0^{-1}$ is shown. These figures outline the cautious approach that `MDiNE` took in determining network edges, and underlines the importance of adequate sample size in proper network inference. The penalization in `MDiNE` pushed all elements of the precision matrices towards zero. We have also shown that there are significant challenges incurred when attempting to perform inference on network edges even for a moderate number of taxa.

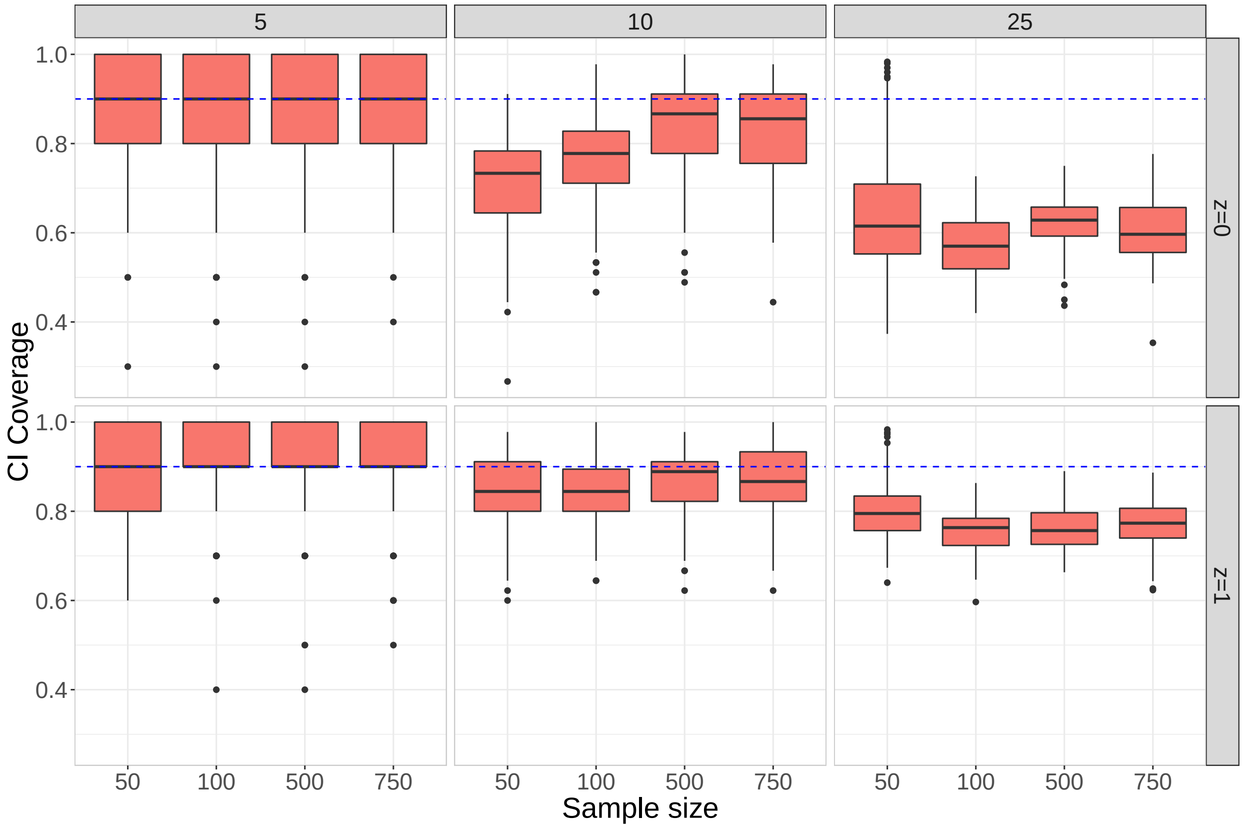


Figure S7: The 90% credible interval coverage averaged over the different elements of $\mathbf{\Sigma}_0^{-1}$ and $\mathbf{\Sigma}_1^{-1}$. The boxplots show the distribution of the average credible interval coverage over the simulation replications.

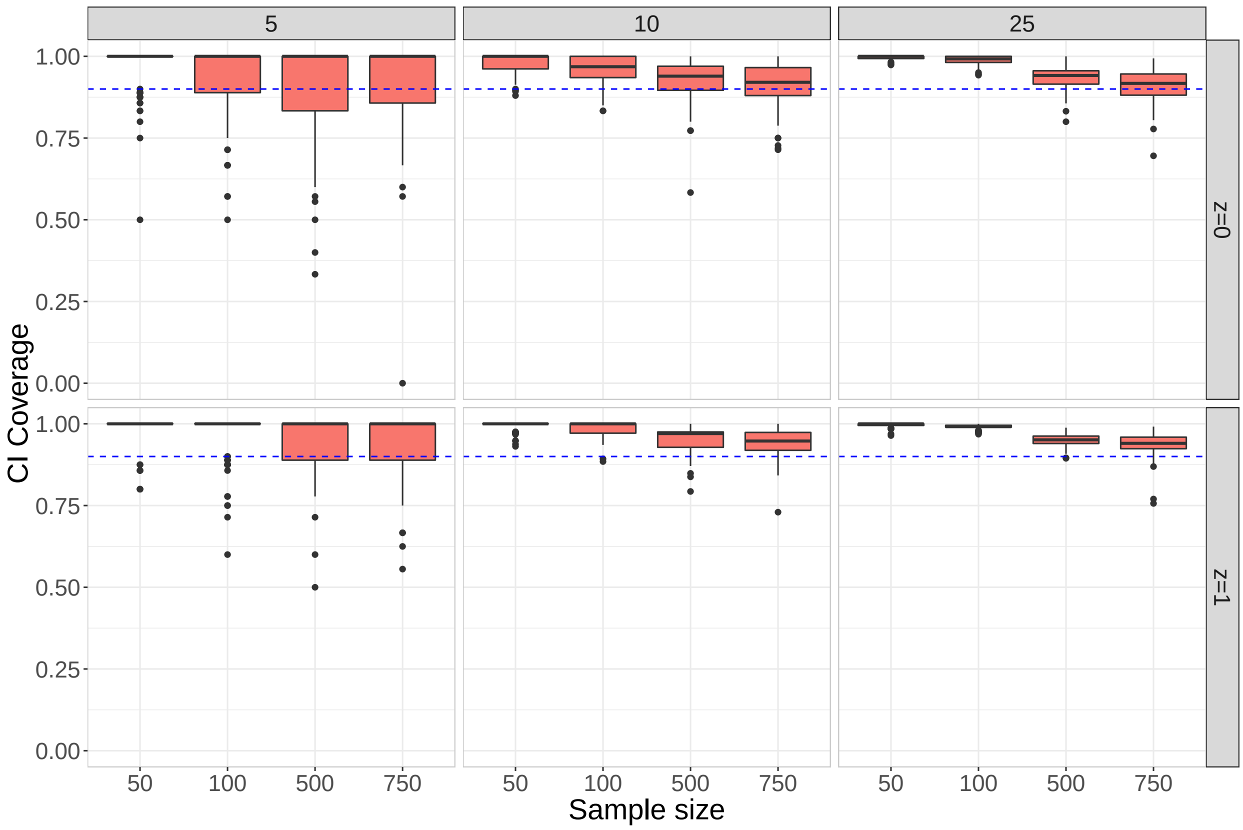


Figure S8: The 90% credible interval coverage averaged over the **true zero** elements of $\mathbf{\Sigma}_0^{-1}$ and $\mathbf{\Sigma}_1^{-1}$. The boxplots show the distribution of the average credible interval coverage over the simulation replications.

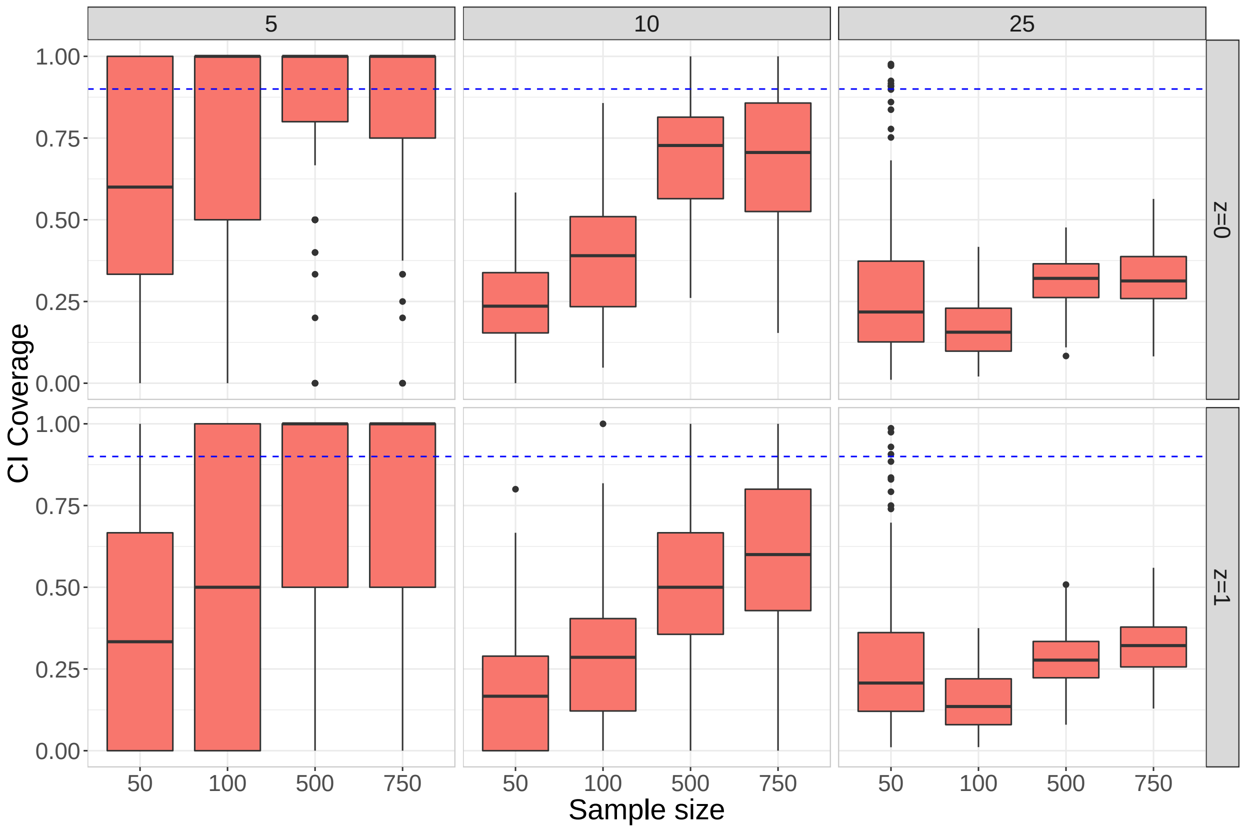


Figure S9: The 90% credible interval coverage averaged over the **true non-zero** elements of $\mathbf{\Sigma}_0^{-1}$ and $\mathbf{\Sigma}_1^{-1}$. The boxplots show the distribution of the average credible interval coverage over the simulation replications.

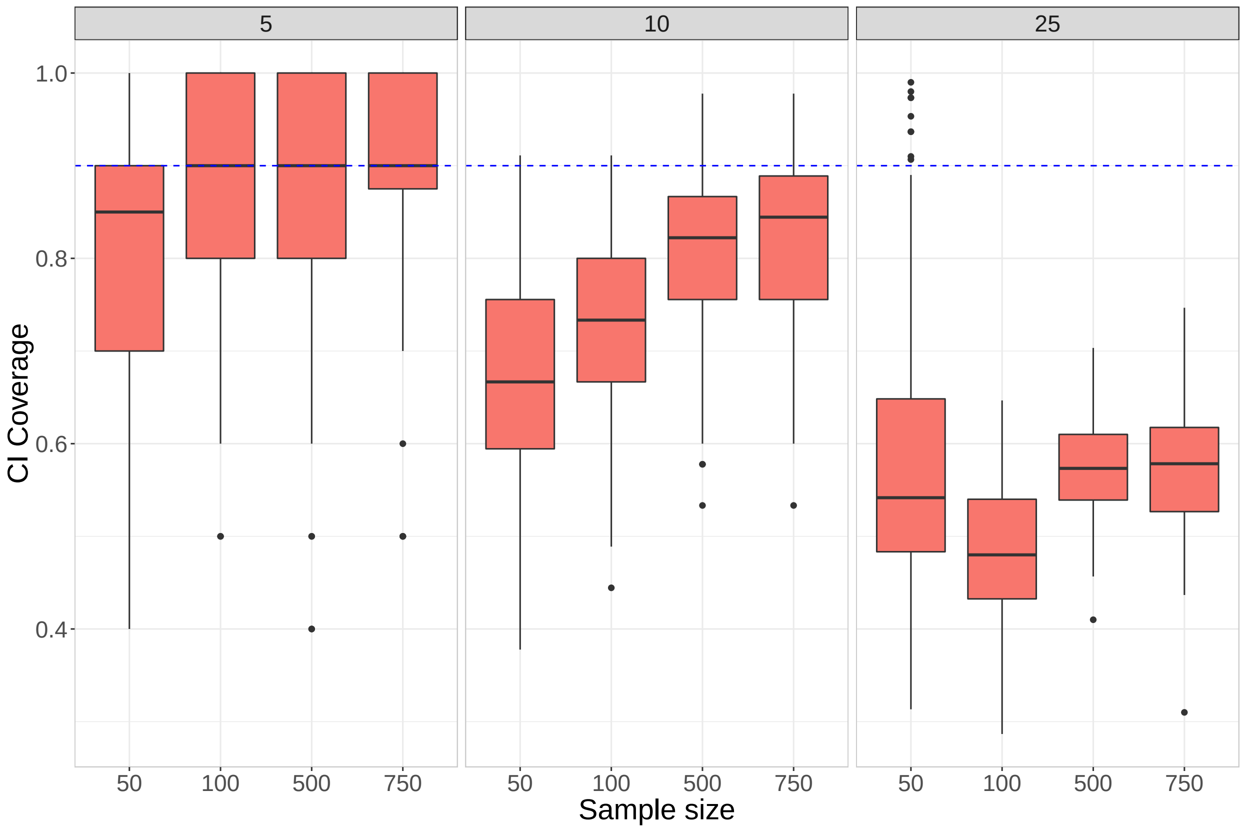


Figure S10: The 90% credible interval coverage averaged over the elements of the difference $\mathbf{\Sigma}_1^{-1} - \mathbf{\Sigma}_0^{-1}$. The boxplots show the distribution of the average credible interval coverage over the simulation replications.

### 7 Crohn's data application: supplementary material

#### 7.1 Comparing results from the network estimation methods

Here we compare the estimated networks from each of `MDiNE`, `SPIEC-EASI`, and `MInt` for the Crohn's data application. In the main manuscript, the networks were estimated for `MDiNE` after including age, sex, and antibiotic status in the design matrix. As `SPIEC-EASI` cannot handle covariates in its model formulation, we ran all three methods assuming no adjustment covariates.

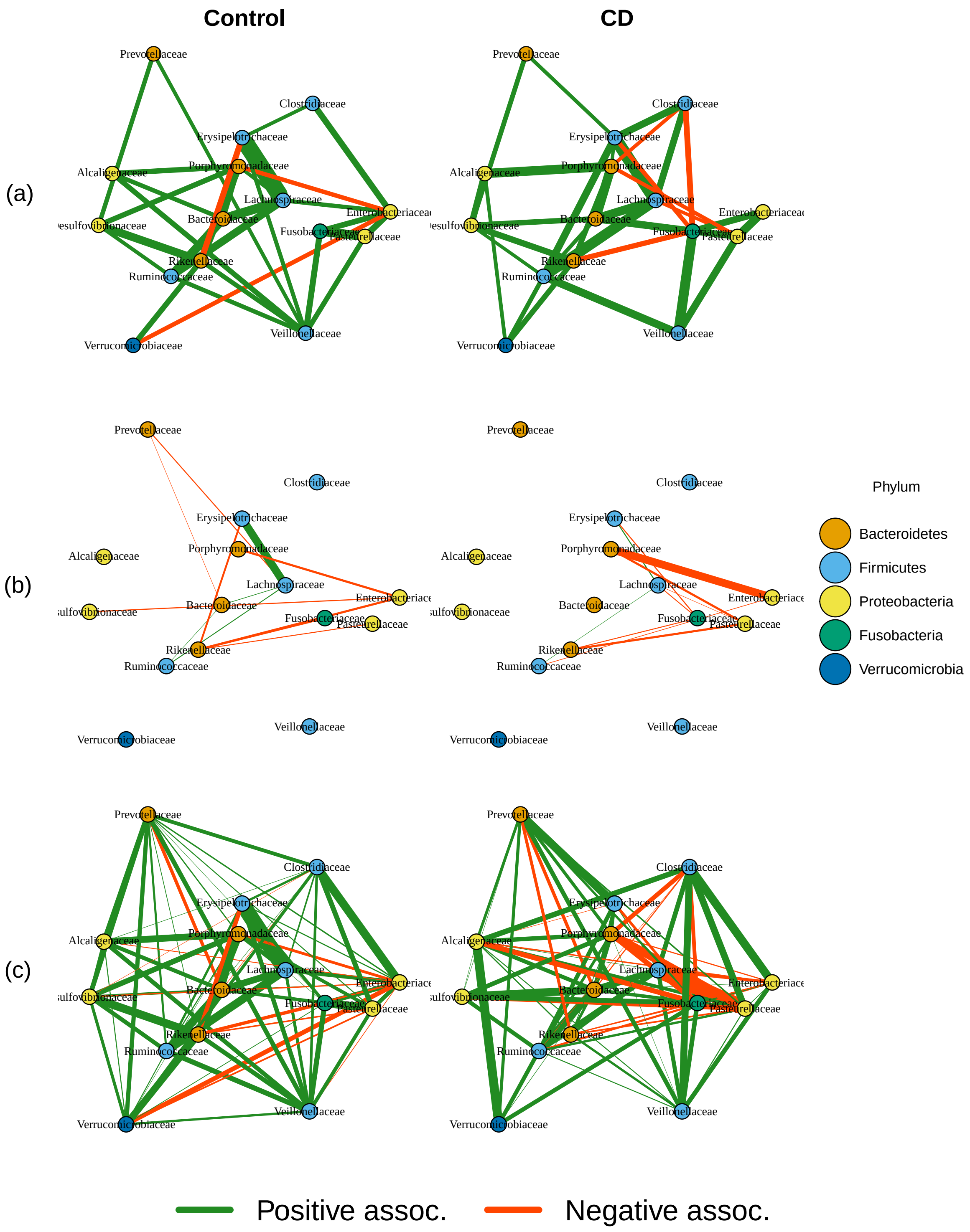


Figure S11: Estimated networks in controls (left) and Crohn's cases (right) from each of the methods: (a) `MDiNE`, (b) `SPIEC-EASI`, and (c) `MInt`. Edge width represents the strength of association.

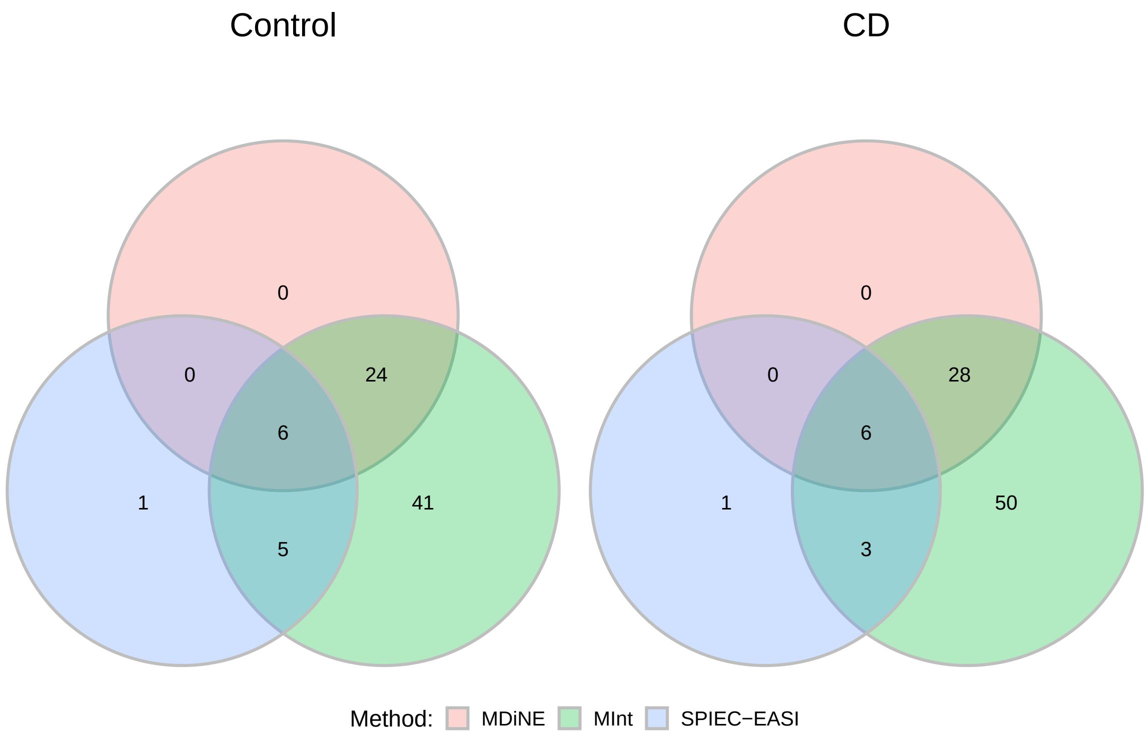


Figure S12: Edge detection concordance between `MDiNE`, `SPIEC-EASI`, and `MInt` in the Crohn's dataset application.

#### 7.2 `MDiNE` convergence diagnostics in Crohn's data application

In order to show that the behaviour of the MCMC sampler was sound, we examined some convergence diagnostics of the analysis of the Crohn's dataset from the main manuscript. First, we examined traceplots to see whether the four chains in each model mixed successfully. Figures S13 and S14 are traceplots for $\widehat{\boldsymbol{\Sigma}}_0^{-1}$ and $\widehat{\mathbf{B}}$, respectively. Mixing was satisfactory for all parameters shown.

We also examined the potential scale reduction factor, often referred to as $\widehat{R}$ (Gelman et al., 1992). This factor compares the amount of between-chain variation with the amount of within-chain variation. If $\widehat{R}$ is much larger than 1, then the chains have not yet adequately mixed, and the model would benefit from additional sampling. If $\widehat{R}$ is approximately 1, then no additional sampling is needed. $\widehat{R} < 1.2$ has been suggested as a reasonable cut-off to determine whether the chains have converged (Brooks and Gelman, 1998).

Figures S15 and S16 show the value of $\widehat{R}$ for each element of $\widehat{\boldsymbol{\Sigma}}_0^{-1}$ and $\widehat{\mathbf{B}}$, respectively. The value of the potential scale reduction factor was very close to 1 in almost all cases. There is clear evidence that the parameters have converged in the Crohn's network analysis presented in the main text.

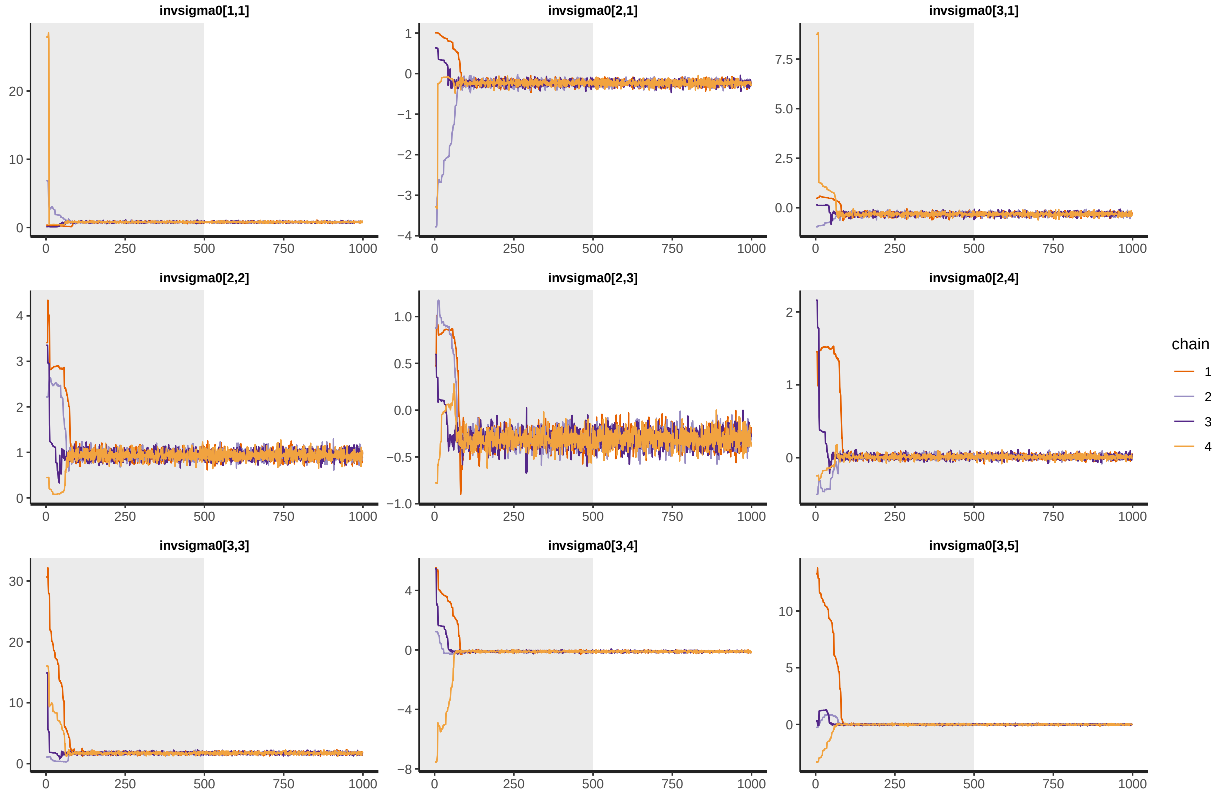


Figure S13: Traceplots of nine of the elements from $\widehat{\boldsymbol{\Sigma}}_0^{-1}$. The shaded area corresponds to the burn-in period.

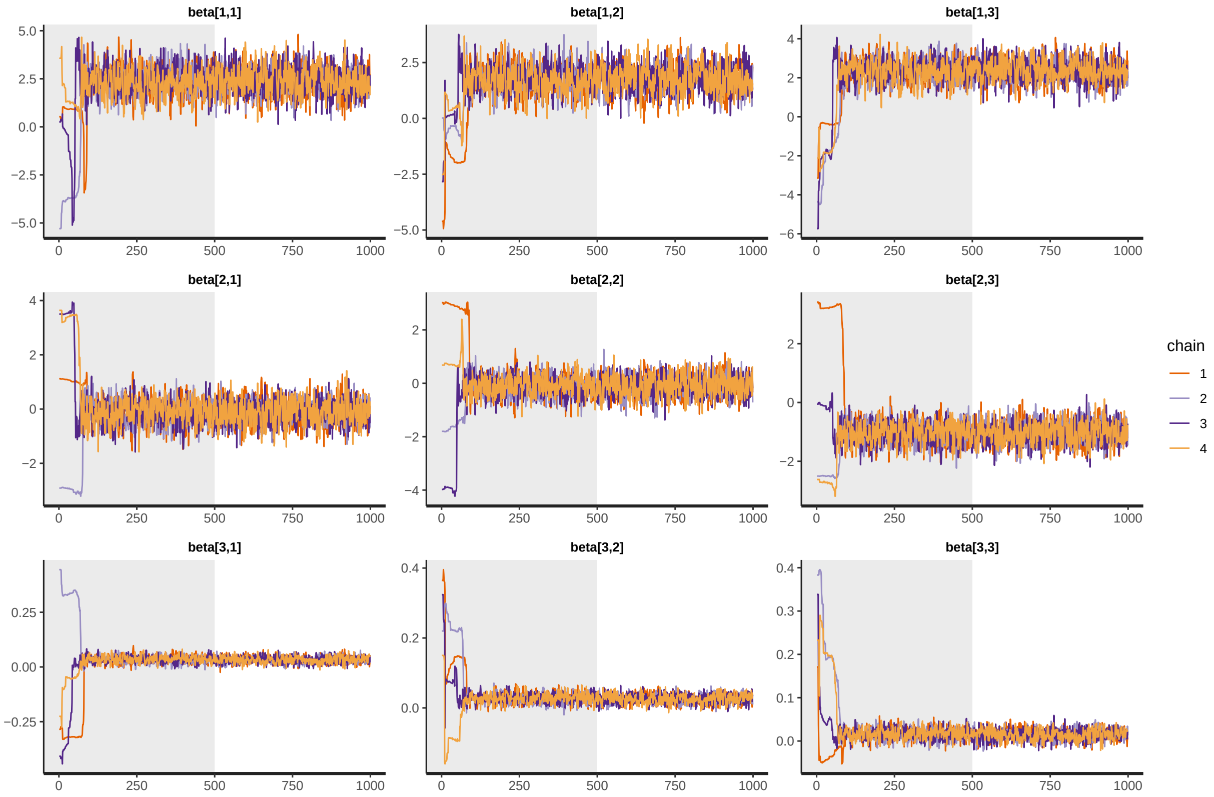


Figure S14: Traceplots of nine of the elements from $\widehat{\mathbf{B}}$. The shaded area corresponds to the burn-in period.

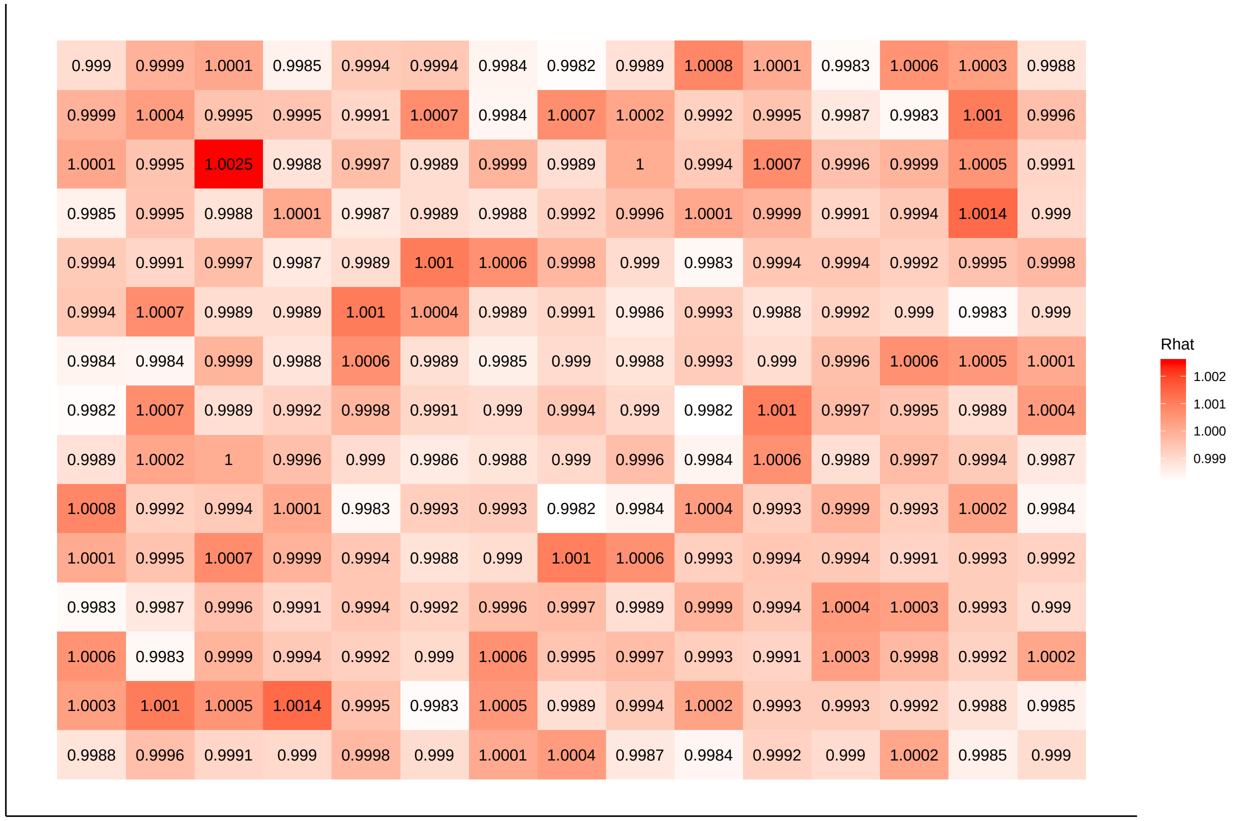


Figure S15: The potential scale reduction ($\widehat{R}$) of the individual elements of $\widehat{\boldsymbol{\Sigma}}_0^{-1}$ in the Crohn's data application.

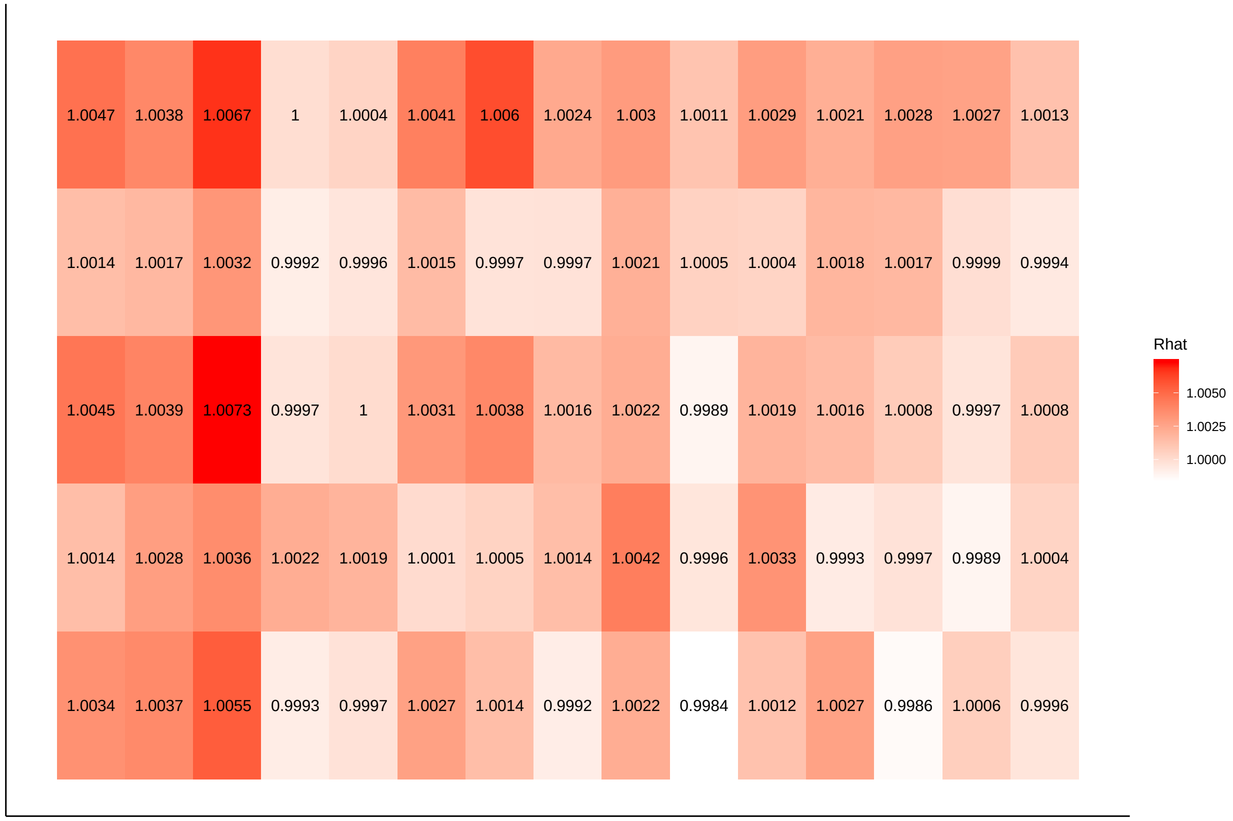


Figure S16: The potential scale reduction ($\widehat{R}$) of the individual elements of $\mathbf{B}$ in the Crohn's data application.

Rui Fang. *Zero-inflated negative binomial (ZINB) regression model for over-dispersed count data with excess zeros and repeated measures, an application to human microbiota sequence data*. PhD thesis, University of Colorado at Denver, Anschutz Medical Campus. Health Sciences . . . , 2013.

Andrew Gelman, Donald B Rubin, et al. Inference from iterative simulation using multiple sequences. *Statistical science*, 7(4):457–472, 1992.

Paul J McMurdie and Susan Holmes. phyloseq: an r package for reproducible interactive analysis and graphics of microbiome census data. *PloS one*, 8(4):e61217, 2013.

Brian Ripley, William Venables, and Maintainer Brian Ripley. Package ‘nnet’. *R package version*, pages 7–3, 2016.

Zhang Xiao-Ke, Wu Jun, Tan Yue-Jin, Deng Hong-Zhong, and Li Yong. Structural robustness of weighted complex networks based on natural connectivity. *Chinese Physics Letters*, 30(10):108901, 2013.
